# Cellular origin and therapeutic potential of SERPINA3N in ischemic stroke

**DOI:** 10.64898/2026.07.27.741013

**Authors:** Joohyun Park, Hyun Kyu Kim, Bhabotosh Barman, Raymond A. Sobel, Midori A. Yenari, Fuzheng Guo

## Abstract

Murine serine protease inhibitor clade A member 3N (SERPINA3N) and its human homolog SERPINA3 are injury-associated molecules induced in virtually all neurological conditions including ischemic stroke. Clinically, SERPINA3 is elevated in stroke patients, correlating with severe brain pathology and worse neurological outcomes. Yet, the exact cellular sources and therapeutic value of SERPINA3N/SERPINA3 in the ischemic brain remain elusive. Here, we report that oligodendrocytes are the major sources of SERPINA3/SERPINA3N and that SERPINA3N exerts neurotoxic effects in experimental cerebral ischemia. Given SERPINA3N’s secretory nature, we used knock-in Serpina3n-tdTom reporter mice to define its temporospatial evolution. The tdTom signal was barely detectable by day1 but gradually increased from day 3 through day 7 following cerebral ischemia. The majority of tdTom^+^ cells were identified as oligodendrocytes and astrocytes with fewer found in neurons or microglia. This cellular identity was corroborated by meta-analysis of single-cell transcriptomics datasets and human ischemic stroke brain data. Antagonizing SERPINA3N attenuated blood-brain barrier disruption, neuroinflammation, and neuronal damage and improved neurological function in a mouse model of cerebral ischemia. These findings resolve a longstanding debate regarding the cellular origins of SERPINA3N, identifying oligodendrocytes as its major source. Our studies highlight SERPINA3N antagonism as a possible therapeutic intervention for ischemic stroke, providing new insights for precision therapies that target oligodendroglia- or astroglia-specific SERPINA3N.

## Introduction

Ischemic stroke is defined as an interruption of blood supply to the brain. The goal of therapy for acute ischemic stroke is to restore blood flow^1^, which unfortunately can lead to secondary reperfusion-oxidative damage to the brain through robust molecular dysregulation and cellular dysfunction^2^. There is no cure for brain secondary injury primarily because of our incomplete understanding of the underlying molecular mechanisms.

Serine protease inhibitor clade A member 3N (SERPINA3N) and its human orthologue SERPINA3^3^ are acute-phase glycoproteins predominantly secreted by the liver in response to systemic inflammation^4^. SERPINA3/SERPINA3N is upregulated across almost all neurological conditions and animal models^4^, such as multiple sclerosis^5^, Alzheimer disease^6,7^, normal aging^8^, neurotrauma^9^, and stroke^10,11^. However, the exact cellular origins of SERPINA3/SERPINA3N in the ischemic brain is highly debated and yet to be resolved. A 2012 study reported that *Serpina3n* mRNA transcript was upregulated almost exclusively in reactive astrocytes in the murine brain subjected to transient middle carotid artery occlusion (tMCAO), a widely used model for studying ischemic stroke pathogenesis^12^. In sharp contrast, recent studies reported that SERPINA3N was expressed predominantly in neurons^10,13^ and, to a lesser extent, in astrocytes following experimental ischemic stroke^10^. Considering SERPINA3N’s secretory properties, SERPINA3N immunoreactive signal experimentally detected in one cell type may come from other cell types. Hence, it is important and necessary to use genetic reporter mice to define the cellular sources of SERPINA3N in the brain following cerebral ischemia.

The functional significance of aberrant SERPINA3/SERPINA3N expression remains enigmatic. Clinically, SERPINA3N was reported to be elevated in stroke patients^14^. The elevated plasma level of SERPINA3 correlated with poor neurological outcomes in patients with hemorrhagic stroke ^11,14^, suggesting that SERPINA3N may play a neurotoxic role in stroke pathophysiology. A recent preclinical study reported that astrocyte-derived SERPINA3N exerted neuroprotective effects by attenuating neuroinflammation in a mouse model of ischemic stroke ^10^. More recently, neuron-derived SERPINA3N was reported to act as an endogenous protector of the blood-brain barrier (BBB) in mice following cerebral ischemic stroke ^13^. Although the growing body of evidence suggests that SERPINA3N is dynamically regulated in ischemic stroke, its precise cellular origins and functional contributions to ischemic brain remain incompletely understood. These knowledge gaps make it necessary to determine whether SERPINA3N/SERPINA3 induction in the brain is a maladaptive injury perpetuator or a compensatory protective factor of post-ischemic responses and injuries. Answers to these questions will provide novel insights into clinical trials that intervene in SERPINA3/SERPINA3N to mitigate ischemic stroke pathology.

In the present study, we aimed to define the cellular origins of SERPINA3N and decipher the therapeutic value of SERPINA3N in post-stroke brain pathology in mice exposed to tMCAO. Unexpectedly, we found that oligodendrocytes are the major cell population expressing SERPINA3N and in contrast to previous reports, neurons exhibited limited SERPINA3N induction. Antagonizing SERINA3N after tMCAO attenuates ischemic brain pathology including BBB dysfunction and neuroinflammation, ultimately alleviating neurological function impairment. Our findings suggest that SERPINA3N induction is a maladaptive mechanism that potentiates post-stroke brain injury.

## Results

### SERPINA3N induction shows temporal and spatial evolution in the tMCAO mouse model of cerebral ischemia

To characterize endogenous expression of the secretory protein SERPINA3N following cerebral ischemia, we performed transient middle cerebral artery occlusion (tMCAO) surgery (sham surgery as control) to *Serpina3n-P2A-tdTomato* reporter mice generated in our recent study ^15^. In Serpina3n-tdTom mice, the fusion protein Serpina3n-tdTom is expressed under the control of endogenous mouse SERPINA3N protein translation. The self-cleavage site P2A ensures separation of tdTom from SERPINA3N and the tdTom reporter remains in SERPINA3N- producing cells ^15^, thus providing a reliable genetic tool to define the temporal and spatial evolution of SERPINA3N expression.

In sham control mice, no tdTom was observed in the brain regions, for example, cerebral cortex (CTX), subcortical white matter (WM), and striatum (STR) (**Fig. 1A**). In tMCAO mice, tdTom expression exhibited a gradual temporal and spatial evolution in the brain following cerebral ischemic. Limited expression of tdTom was observed during the acute stage by day 1 following cerebral ischemia, although massive neuronal damage occurs throughout the ischemic lesion as indicated by the prominent loss of neuronal marker MAP2a (**Fig. 1B**). tdTom-expressing cells increased in quantity from day 3 through day 7 post-stroke (**Fig. 1E**) specifically located in the lesional areas of the ipsilateral hemisphere, such as CTX, WM, and STR (**Fig. 1C-D**). In contrast, minimal tdTom expression was detected in the contralateral hemisphere of tMCAO mice (**Fig. 1C-E**). These data are in line with our recent study showing that SERPINA3N is an injury-induced molecule that is otherwise absent from the homeostatic brain ^15^. The temporal regulation was corroborated by Western blot analysis showing that endogenous SERPINA3N gradually increased from day 3 to day 7 post-stroke (**Fig. 1F**). Collectively, these findings demonstrate that SERPINA3N undergoes delayed and lesion-associated induction following cerebral ischemia, with maximal expression observed after the acute phase post-stroke.

**Figure 1.**
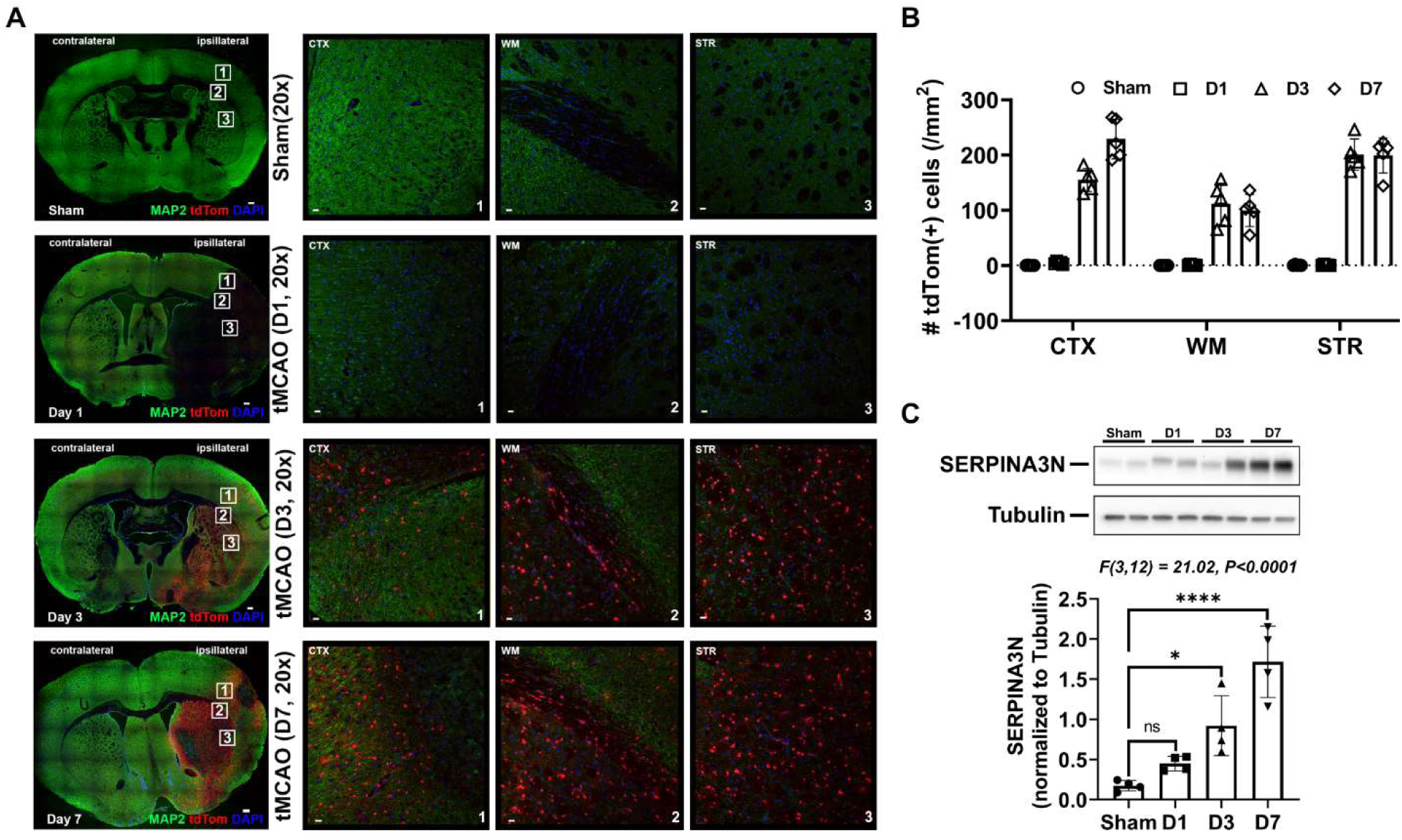
- Temporal and spatial evolution of Serpina3n-expressing cells during cerebral ischemia. (A) Representative immunofluorescence images showing the temporal and regional distribution of tdTom (as a reporter of Seprina3n transcription) in the cortex (CTX), white matter (WM), and striatum (STR) at indicated timepoints after cerebral ischemia. Loss of the neuronal marker MAP2 was used to indicate neurodegeneration. Boxed areas indicate higher magnification views of the corresponding numbered regions (1, CTX; 2, WM; 3, STR). Scale bar, 20 µm. (B) Quantification of tdTom+ cells in each ipsilateral region over time following cerebral ischemia. (C) Representative immunoblot and quantification of SERPINA3N protein expression over time following cerebral ischemia. (One-way ANOVA with Tukey’s multiple comparisons test; F(DFn, DFd) and p value; F(3, 12) = 21.02, p<0.0001, ns, not significant; n=4 per group)

### The identity of SERPINA3N-expressing cells shifts in a time-dependent manner after cerebral ischemia

To identify the cellular sources of upregulated SERPINA3N protein in the ischemic brain, we performed IHC of tdTom and lineage-specific markers. We used NeuN for identifying neurons, GFAP for astrocytes, GSP-pi for oligodendrocytes, and CD68 for activated myeloid cells including microglia/macrophages. We took the cerebral cortex as a representative for brain cortical regions and corpus callosum for subcortical white matter regions. As expected, we did not find consistent tdTom expression in cells located at the contralateral hemisphere to the ischemic stroke injury, for instance NeuN^+^ neurons and GST-pi^+^ oligodendrocytes (**Suppl Fig. 1A-B**). CD68 and GFAP expression was minimal in the contralateral hemisphere throughout the disease course (**Supple Fig. 1C-D**) presumably reflecting the limited injury and glial activation. In the ipsilateral hemisphere, prominent CD68 expression was absent by day 1 (**Suppl Fig. 2A- B**) and robust by day 3 post-stroke injury, but it was mutually exclusive from tdTom (**Suppl Fig. 2C-D)**, suggesting that activated microglia and macrophages are not a cellular source of SERPINA3N in the ischemic brain.

Double IHC of tdTom and NeuN demonstrated that tdTom expression was absent from NeuN^+^ neurons from day1 through day 7 we assessed (**Fig. 2A, E**). Interestingly, double IHC of SERPINA3N and NeuN indicated that SERPINA3N immunoreactive signal appeared to colocalize with NeuN^+^ neurons at the acute phase by day 1 (**Suppl Fig. 3A**). However, IHC on the adjacent cryosections clearly showed the absence of tdTom from NeuN^+^ neurons (**Suppl Fig. 3B**). We found that almost all tdTom^+^ cells were identified as GST-pi^+^ oligodendrocytes and GFAP^+^ astrocytes in the cerebral cortex (**Fig. 2B-C**) and corpus callosum (**Fig. 2F-G**) of ischemic brain. In the cerebral cortex, ∼60% of tdTom^+^ cells were identified as astrocytes at day 3 and this proportion decreased to less than 10% by day 7. Conversely, ∼40% of tdTom+ cells were identified as oligodendrocytes at day 3 and this proportion increased over 90% by day 7 post-stroke (**Fig. 2D**). In the corpus callosum, over 70% of tdTom+ cells were identified as oligodendrocytes throughout the time course with less than 10% being astrocytes (**Fig. 2H**). These data demonstrate a temporal change in SERPINA3N-expressing cell identities during the time course of post-stroke secondary injury. Taken together, these findings reveal a previously underappreciated oligodendroglia-dominant and temporally regulated pattern of SERPINA3N expression following ischemic stroke. Contrary to the prevailing neuron-centric view, our genetic reporter data convincingly identify oligodendrocytes, and to a lesser extent reactive astrocytes, as the predominant SERPINA3N-expressing cell populations in the ischemic brain.

**Figure 2.**
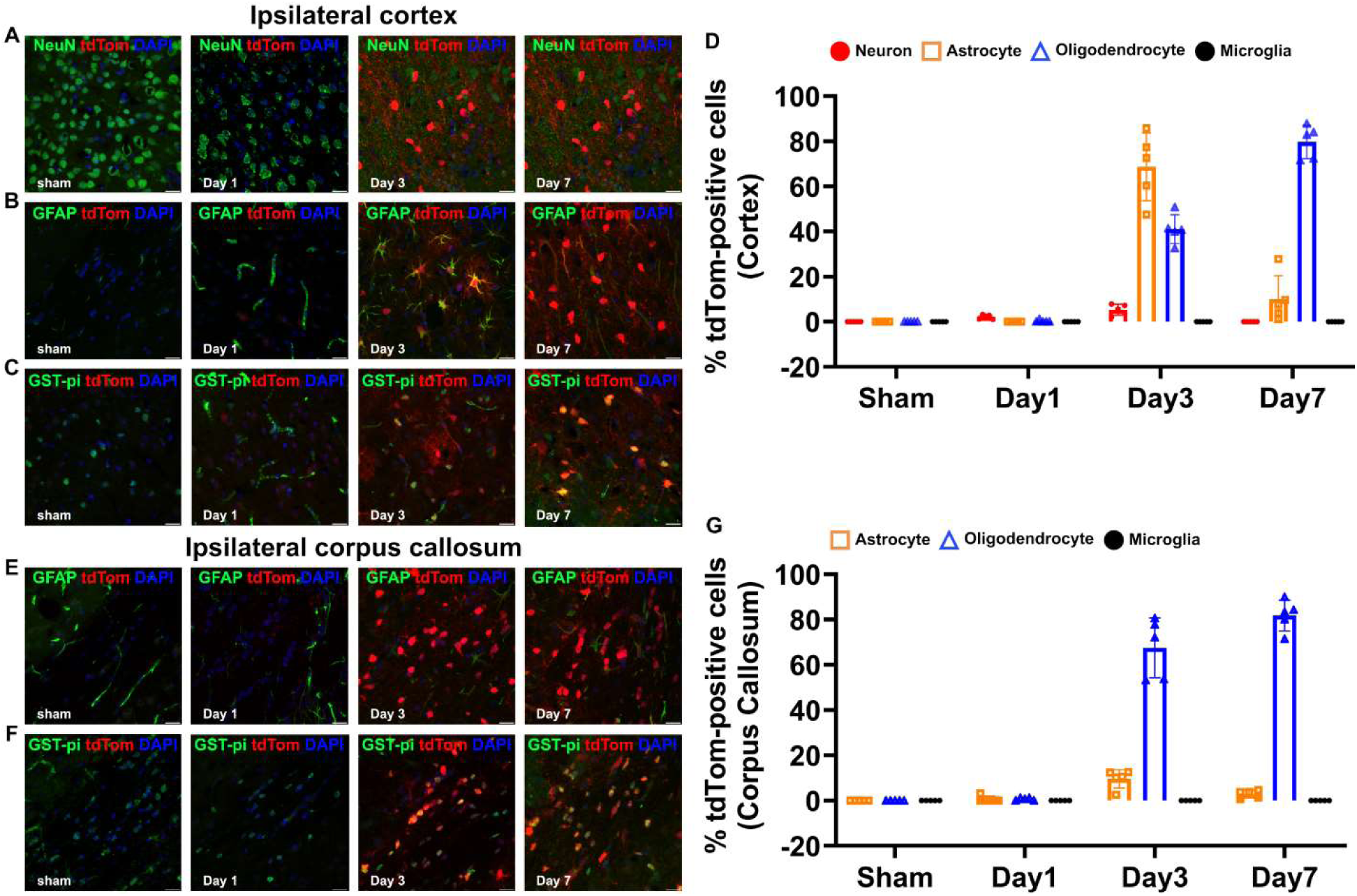
– Cellular identity of Serpina3n-expressing cells during cerebral ischemia. **(A-C)** Representative images of tdTom and lineage-specific markers (NeuN, neurons; GFAP, astrocytes; GSP-pi, oligodendrocytes) in the CTX and **(E, F)** CC at indicated timepoints following cerebral ischemia. Scale bars, 20 µm. n=5 per group. **(D, G)** Percentage of tdTom+ cells that are positive for lineage-specific markers in the cortex and corpus callosum at different time points following cerebral ischemia. Mean ± SD.

### Meta-analysis of single cell transcriptomics datasets reveals glial upregulation of Serpina3n mRNA

To strengthen our conclusion that oligodendrocytes but not neurons are the major cellular source of SERPNA3N protein in the ischemic brain, we sought to analyze single cell RNA-seq of brain cells after tMCAO. The datasets we used were publicly deposited by three independent laboratories^16–18^, thus avoiding the potential influence of surgery, sampling, and technical differences across datasets on data interpretation. After data integration and quality control, we identified 14 different cell populations including glial cells, neurons, immune myeloid and lymphoid cells, vascular cells, and fibroblasts (**Fig. 3B**) based on the marker gene expression (**Fig. 3A**). Our results demonstrated that the highest upregulation of Serpina3n mRNA was in oligodendrocytes, and to a lesser extent, in astrocytes (**Fig. 3C**). Conversely, neurons and microglia exhibited low levels of Serpina3n mRNA expression (**Fig. 3C**). We found that among the Serpin clade A (Serpina) members, Serpina3n mRNA displayed the highest level of induction by the occlusion-reperfusion injury after tMCAO compared to sham group (**Fig. 3D**). The RNA-seq results confirmed our conclusion drawn from Serpina3n-tdTom reporter mice, demonstrating that oligodendrocytes, but not neurons, are the major cell type producing SERPINA3N post-stroke.

**Figure 3.**
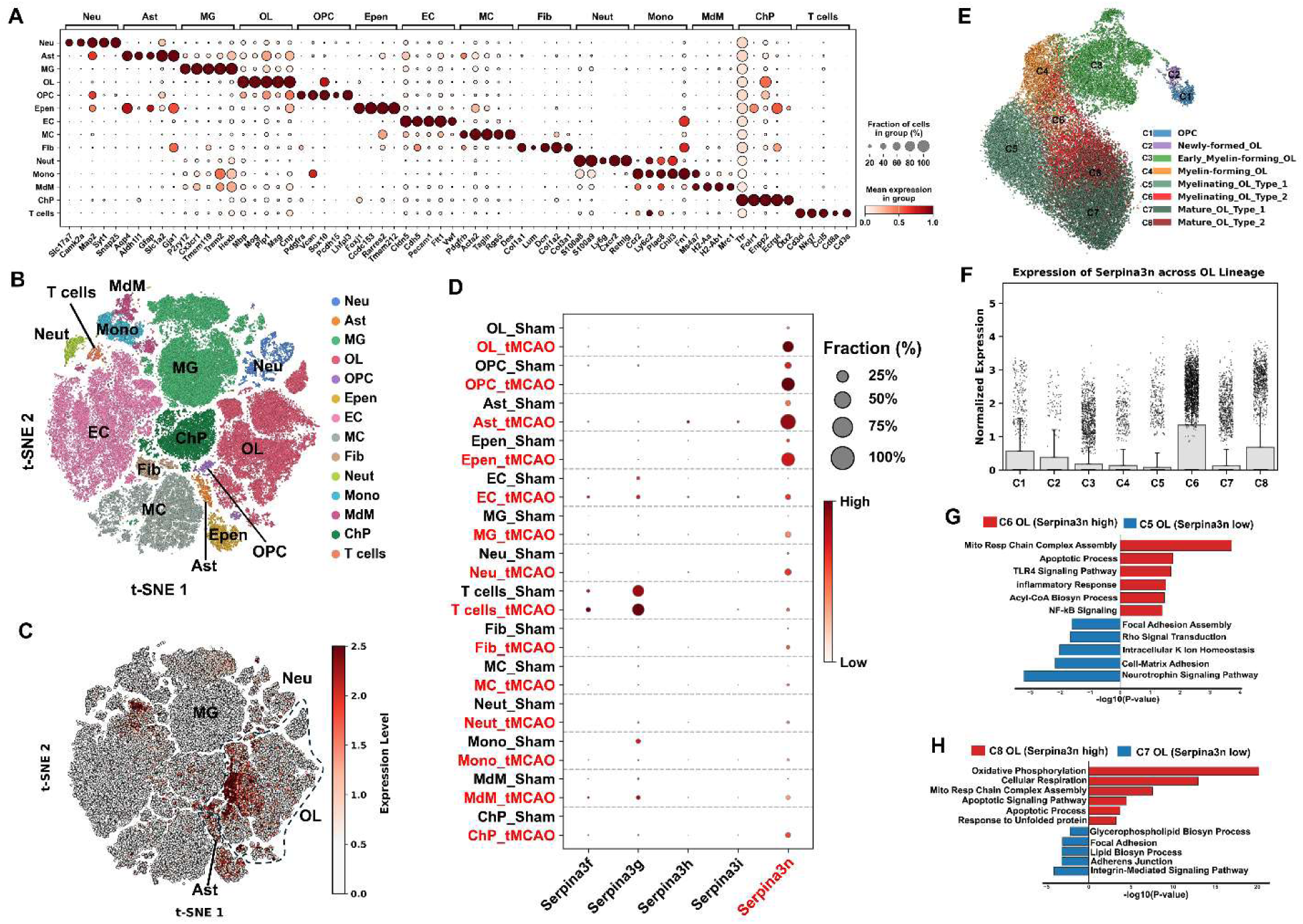
–single-cell transcriptomic meta-analysis identifies oligodendrocytes as the major cellular source of Serpina3n in the tMCAO model. **(A)** Dot plot illustrating the expression profiles of canonical marker genes across 14 distinct cell populations: neurons (Neu), astrocytes (Ast), microglia (MG), oligodendrocytes (OL), oligodendrocyte precursor cells (OPC), ependymal cells (Epen), endothelial cells (EC), vascular mural ells (MC), fibroblasts (Fib), neutrophils (Neut), monocytes (Mono), monocyte-derived macrophages (MdM), choroid plexus cells (ChP), and T cells. Node size represents the fraction of cells expressing the target gene within a specific cluster, while color intensity denotes the mean normalized expression level. **(B)** t-SNE projection of the combined meta-dataset from three independent tMCAO cohorts, color-coded by the 14 annotated cell types. **(C)** Feature plot showing the relative levels of Serpina3n mRNA across cell populations. **(D)** Dot plot graph showing the percentage and relative level of Serpina3n and other SerpinA family members in each cell population between sham and tMCAO groups. **(E)** Sub-clustering of oligodendrocyte lineage cells. **(F)** Relative expression of Serpina3n mRNA in each subcluster cell. Note that Serpina3n mRNA was upregulated in C6 myelinating OLs type 2 (versus C5 myelinating OLs type 1) and C8 mature OLs type 2 (versus C7 mature OLs type 1). **(G)** Functional enrichment in myelinating OL subpopulations (C6 vs. C5). Bar chart detailing the significantly enriched biological pathways in the Serpina3n-high Type 2 phenotype (C6, red) versus the Serpina3n- low Type 1 phenotype (C5, blue). **(H)** Major biological processes enriched in type 2 versus type 1 mature OLs. Three datasets of single cell RNA-seq from independent laboratories were included in the meta-analysis: GSE19773118, GSE22765117, GSE24538616.

Next, oligodendroglial lineage cells were further subclustered into different developmental and injury stages from OPCs through mature OLs (C1-C8) (**Fig. 3E**). We found that Serpina3n mRNA was dramatically upregulated in myelinating OLs type 2 (versus myelinating OLs type 1) and mature OLs type 2 (versus mature OLs type 1) (**Fig. 3F**), suggesting that type 2 myelinating and mature OLs are the major cell subclusters that respond to occlusion-reperfusion injury after tMCAO. Gene enrichment analysis showed that Serpina3n-dominant type 2 cells (C8 and C6) demonstrated marked enrichment in the biological processes related to mitochondria respiratory chain assembly, apoptotic pathway, and inflammatory response (**Fig.3 G-H**, red). Conversely, type 1 cells of low or no Serpina3n expression (C7 and C5) associated with cellular homeostasis and focal adhesion assembly (**Fig. 3G-H**, bule). These findings indicate that Serpina3n induction may be associated with brain inflammation and cell apoptosis.

### Oligodendrocytes are the major cell types expressing SERPINA3 in the brain of ischemic stroke patients

We next asked whether SERPINA3N expression in tMCAO mice is conserved in the brain of ischemic stroke patients. To this end, we analyzed the post-mortem brain specimens using formalin-fixed paraffin-embedded (FFPE) sections. Using hematoxylin/eosin (HE) staining, we identified ischemic lesion cores and peri-infarct areas on FFPE sections (**Fig. 4A**). Using adjacent sections, we observed that SERPINA3, the human ortholog of murine SERPINA3N was expressed in the peri-infarct regions of cerebral cortex and white matter). The identity of SERPINA3-expressing cells was determined by double IF staining with lineage-specific markers: Neu for neurons, GFAP for astrocytes, GST-pi for oligodendrocytes, and CD68 for activated microglia/macrophages (**Fig. 4B**). In the peri-infarct cortical area, GST-pi^+^ oligodendrocytes represented the predominant SERPINA3-expressing cell population, followed by astrocytes (**Figure 4C**). Similarly, in peri-lesion white matter area, oligodendrocytes constituted the major SERPINA3-positive population, followed by astrocytes (**Figure 4D**). Consistent with the rodent findings, only a very small population of neurons exhibited SERPINA3 immunoreactivity and no detectable colocalization between SERPINA3 and microglia/macrophages was observed (data not shown here). Collectively, these findings demonstrate that SERPINA3 expression in the post-stroke brain is predominantly enriched in oligodendrocytes, with substantial contribution from reactive astrocytes, thereby supporting a glia-centered pattern of SERPINA3 expression following cerebral ischemia.

**Figure 4.**
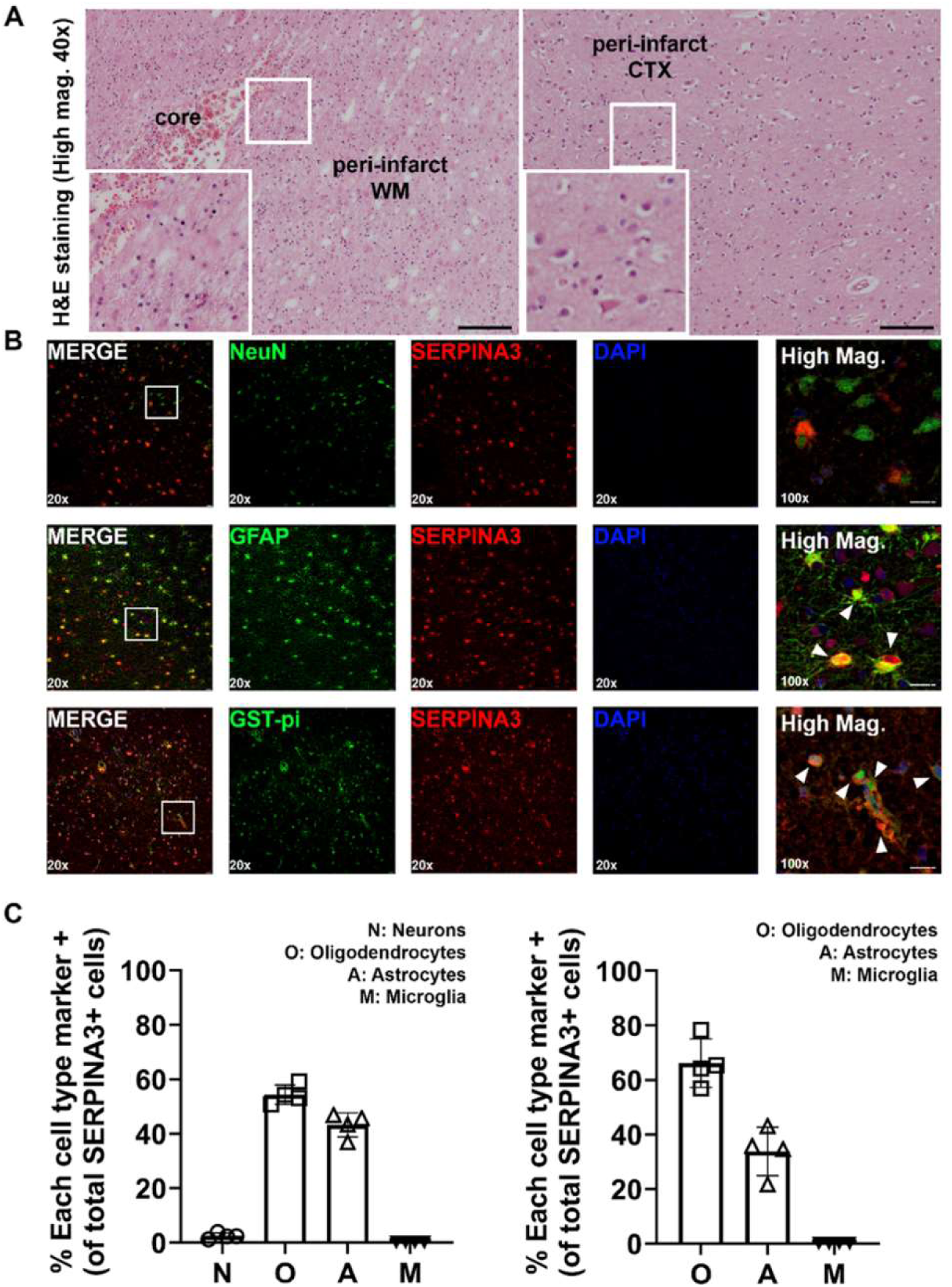
– Oligodendrocytes are the predominant cellular sources of SERPINA3 in human post-stroke brain tissue. **(A)** Representative hematoxylin and eosin (H&E) staining of human post-stroke brain tissue identifying the infarct core, peri-infarct cortex, and white matter regions. White boxed areas indicate regions containing morphologically damaged neurons and oligodendrocytes, which are shown at higher magnification (40x). Scale bar, 1 mm. **(B)** Representative immunofluorescence images of SERPINA3 expression across neural cell populations in brain tissue, co-stained with cell type-specific markers for neurons (NeuN), astrocytes (GFAP), oligodendrocytes (GST-pi), and microglia (data are not shown). White boxed regions are presented at higher magnification (100x). Scale bar, 20 µm. Magnification, 20x. **(C)** Quantitative analysis of SERPINA3+ cells by cell type in in peri-infarct cortex (left) and white matter (right). SERPINA3 expression was predominantly detected in oligodendrocytes (Cortex, 54.34 ± 3.56; WM, 66.18 ± 8.88), to a lesser extent, in astrocytes (Cortex, 43.21 ± 4.45; WM, 33.82 ± 8.88), whereas barely detectable in neurons (Cortex: 2.45 ± 1.09) and no expression was observed in microglia. (n=4 per group).

### Antagonizing SERPINA3N attenuates blood-brain barrier (BBB) impairment and neurological impairment after tMCAO

To investigate the potential contribution of brain SERPINA3N induction to post-ischemic brain pathology, we treated mice with SERPINA3N inhibitor ARN1468^19^ through intraperitoneal injections twice a day right after tMCAO surgery. To study the effect of ARN1468 on BBB functional integrity, we injected Evans blue (EB) 30 minutes prior to animal perfusion and brain harvest, a dye that is widely used to evaluate BBB permeability. We found minimal EB signal in the sham brain and intense signal in the tMCAO brain treated with vehicle (**Fig. 5A**), indicating disrupted BBB integrity in the ischemic brain. Our quantification showed that ARN1468 significantly reduced EB signal in the brain at day 1 post-tMCAO (**Fig. 5B**), suggesting that SERPINA3N inhibition protects against acute BBB breakdown. Tight junction proteins, such as Occludin (Ocln), Claudin 5 (Cldn5), and Zonula Occludens-1 (Tjp1) play essential roles in maintaining BBB functional integrity. Consistent with the EB data, the expression levels of Ocln, Cldn5, and Tjp1 were remarkably increased in the brain of tMCAO mice treated with ARN1468 compared with those treated with DMSO Vehicle at 7 days post-tMCAO (**Fig. 5C**) assessed by RT-qPCR. which was confirmed by histological analysis of Claudin 5 (**Fig. 5D**). Matrix metalloproteinase 9 (MMP9) and intercellular adhesion molecule-1 (ICAM-1) in endothelial cells are key drivers of vascular inflammation and BBB disruption ^20–22^. We found that ARN1468 significantly decreased the expression of Mmp9 and Icam1 in the tMCAO brain (**Fig. 5E**). In line with the preserved BBB function, the extravasation of immune cells, such as CD3^+^ T cells and Ly6C/G^+^ neutrophils, into the brain was significantly diminished in the tMCAO brain treated with ARN1468 versus Vehicle control (**Fig. 5F**). We used modified neurological severity score (mNSS) as a criterion for neurological assessment of tMCAO mice and found significantly improved neurological functions in tMCAO mice treated with ARN1468 at day 5 and day 6 after cerebral ischemic injury (**Fig. 5G**). Together, these data suggest that SERPINA3N promotes BBB disruption in cerebral ischemia.

**Figure 5.**
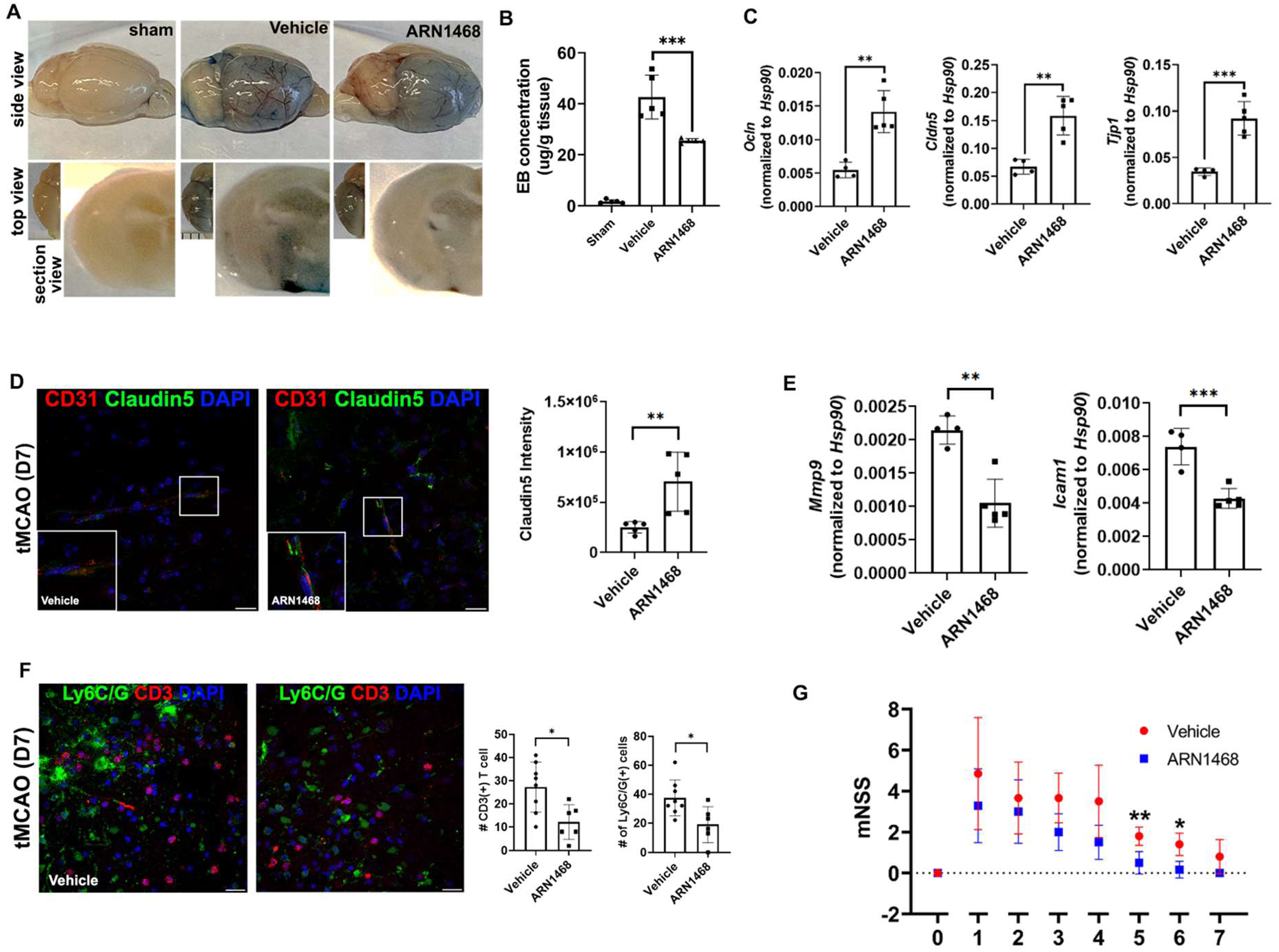
– SERPINA3N inhibition attenuates BBB disruption and neurological severity. **(A-B)** Representative Evans blue staining images and quantification showing BBB breakdown and the effect of ARN1468 at Day 1 post-cerebral ischemia. (One-way ANOVA with Tukey’s multiple comparisons test, F(DFn, DFd), and p value; F(2, 12)=84.39, p<0.0001; n=5 per group) **(C)** RT-qPCR analysis of BBB tight junction gene expression at Day 7 post-cerebral ischemia. (Unpaired Student’s t-test, t(df), and p value; Ocln, t(7)=5.245, p=0.012; Cldn5, t(7)=5.015, p=0.0015; Tjp1, t(7)=6.214, p=0.0004; n=4 Vehicle, n=5 ARN1468) **(D)** Representative immunofluorescence images and quantification of Claudin-5 intensity and quantitative analysis at Day 7 post-cerebral ischemia. (Unpaired Student’s t-test, t(df), and p value; Claudin-5, t(8)=3.366, p=0.0098; n=5 per group) **(E)** RT-qPCR analysis of extracellular matrix degradation (Mmp9) and endothelial adhesion (Icam1) markers of BBB disruption at Day 7 post-cerebral ischemia. (Unpaired Student’s t-test, t(df), and p value; Mmp9, t(7)=5.375, p=0.0010; Icam1, t(7)=5.489, p=0.0009; n=4 Vehicle, n=5 ARN1468) **(F)** Representative immunofluorescence images and quantification of infiltrating immune cells (neutrophils/monocytes, green; T lymphocytes, red; DAPI, blue) at Day 7 post-cerebral ischemia. (Unpaired Student’s t-test, t(df), and p value; Ly6C/G, t(12)=2.740, p=0.0179; CD3, t(12)=2.933, p=0.0125; n=8 Vehicle, n=6 ARN1468) **(G)** Modified neurological severity score (mNSS) in Vehicle and ARN1468-treated mice. Mann-Whitney test (two-tailed); at Day 5, **p=0.0087; at Day 6, *p=0.0108; n=5 Vehicle, n=6

### Antagonizing SERPINA3N mitigates the brain pro-inflammatory response and glial activation after tMCAO

After showing that SERPINA3N promotes BBB disruption, we determined whether SERPINA3N regulates brain inflammatory responses to ischemia. IL-1β (Interleukin-1 beta), TNFα (Tumor necrosis factor alpha), and CXCL12 (C-X-C motif chemokine ligand 12) are all well-established proinflammatory mediators that play critical, destructive roles in BBB and neuronal damage following tMCAO ^23–26^. Our qPCR results showed that the brain levels of IL-1β, TNFα, and CXCL12 were significantly repressed by ARN1468 treatment at day 7 post-tMCAO (**Fig. 6A**), suggesting that SERPINA3N promotes inflammatory response to ischemic injury. We next evaluate the activation of microglia and astroglia which play crucial role in neuroinflammation. Upon activation, microglia downregulates the homeostatic marker and functional genes such Tmem119 and Pyry12^27^. We found that the brain levels of Tmem119 and P2ry12 were significantly increased in the brain of tMCAO mie treated with ARN1468 (**Fig. 6B**). Conversely, ARN1468 treatment decreased the expression of CD68, a pan marker of microglia/macrophage activation (**Fig. 6C**). Similarly, ARN1468 inhibition significantly diminished the brain levels of a panel of pan-markers of reactive astrocytes^28,29^ including GFAP, LCN2, Vim, Cp and Lipg1 (**Fig. 6D**), implying that SEPINA3N promotes astrocyte activation during cerebral ischemia. To corroborate the molecular assays, we used immunofluorescence (IF) to assess glial activation using CD68 and GFAP at the histological level. Double IF staining and quantification showed that the intensity of CD68 and GFAP were significantly decreased in the cerebral cortex (**Fig. 6E**) and striatum (**Fig. 6F**) of tMCAO mice treated with ARN1468 compared with vehicle control group. Collectively, these molecular and histological results imply that SERPINA3N promote neuroinflammation after tMCAO; its antagonism attenuates brain inflammation and microglial/astroglial activation.

**Figure 6.**
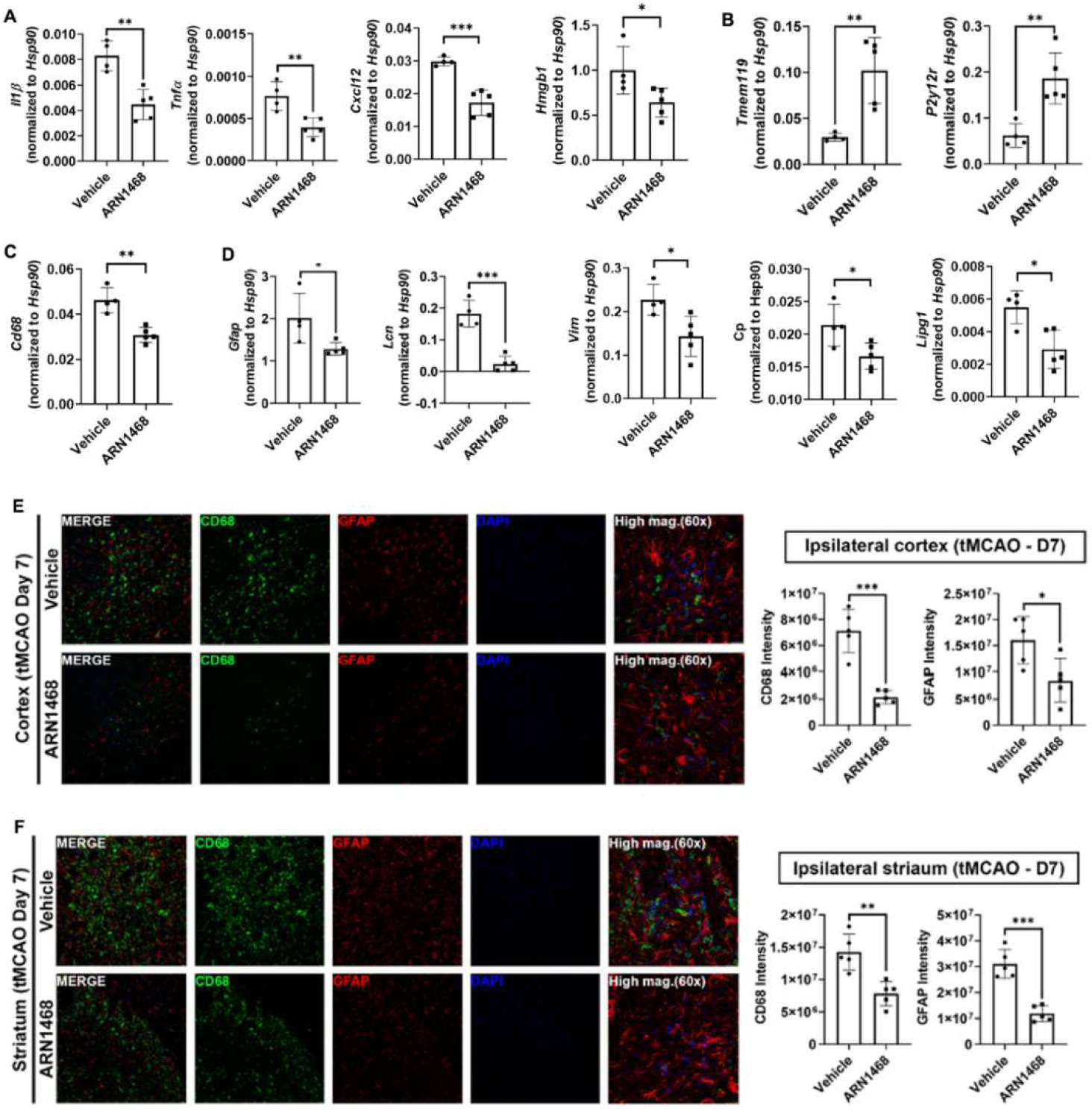
– SERPINA3N inhibition decreases pro-inflammatory response and glial activation in the brain at Day 7 post-cerebral ischemia. **(A)** RT-qPCR analysis of proinflammatory cytokines. (Unpaired Student’s t-test, t(df), and p value; Il1β, t(7)=4.813, p=0.019; Tnfα, t(7)=4.009, p=0.0051; Cxcl12, t(7)=6.163, p=0.0005; Hmgb1, t(7)=2.542, p=0.0386;n= 4 Vehicle, n=5 ARN1468). **(B)** RT-qPCR analysis of microglial homeostatic gene expression. (Unpaired Student’s t-test, t(df), and p value; Tmem119, t(7)=3.923, p=0.0057; P2ry12, t(7)=4.086, p=0.0047; n=4 Vehicle, n=5 ARN1468) **(C)** RT-qPCR analysis of microglial activation gene expression. (Unpaired Student’s t-test, t(df), and p value; Cd68, t(7)=5.142, p=0.0013; n=4 Vehicle, n=5 ARN1468) **(D)** RT-qPCR analysis of reactive astrocytic gene expression. (Unpaired Student’s t-test, t(df), and p value; Gfap, t(7)=2.715, p=0.0030; Lcn, t(7)=7.083, p=0.0002; Vim, t(7)=2.986, p=0.0203; Lipg1, t(7)=3.465, p=0.0105; n=4 Vehicle, n=5 ARN1468) **(E-F)** Representative immunofluorescence images microglia/macrophages (CD68+) and reactive astrocytes (GFAP+) in the cortex and striatum at Day 7. Merged and high magnification images shown. Scale bar, 20 µm. 20x magnification. Quantification of CD68 and GFAP immunoreactivity — cortex: unpaired Student’s t-test, t(df), and p value; CD68, t(8)=6.289, p=0.0002; GFAP, t(8)=2.760, p=0.0247; n=5 per group. Striatum: CD68, t(8)=4.287, p=0.0027; GFAP, t(8)=6.766, p=0.0001; n=5 per group.

### Antagonizing SERPINA3N reduces neuronal damage and death following cerebral ischemia

Neuronal damage and death are the direct substrate for neurological function impairment. We hypothesize that SERPINA3N aggravates neuronal/axonal damage after tMCAO. It is well-established that oxidative stress is a primary driver of neuronal damage in cerebral ischemia^30–33^. We used the monoclonal antibody (clone #15A3), which specifically recognizes markers of oxidative damage to DNA/RNA such as 8-hydroxyguanine, 8-hydroxyguanosine, and 8-hydroxy- 2’-deoxyguanosine, to assess neuronal oxidative stress. As expected, cerebral ischemic injury caused dramatic oxidative stress damage to neurons as evidenced by extensive colocalization of DNA/RNA damage markers and NeuN in the cortex of tMCAO mice (**Fig. 7A**). In sharp contrast, the intensity of DNA/RNA damage markers was significantly diminished in the tMOCA mice treated with ARN1468 (**Fig. 7B**). Oxidative stress damage rapidly induces the expression of immediate-early gene c-Fos in neurons^34^ Consistently, our IF staining (**Fig. 7C**) and quantification showed that the expression of C-Fos was significantly reduced in ARN1468- treated tMCAO mice compared to vehicle control cohort (**Fig. 7D**), which is confirmed by RT-qPCR assay of c-Fos mRNA levels in the brain (**Fig. 7E**). It is well appreciated that neuronal death is primarily through caspase 3-mediated apoptosis during cerebral ischemia ^35,36^. We used cleaved caspase 3 (CC3), the primary executioner of apoptosis to evaluate neuronal death (**Fig. 7F**) and found that ARN1468 treatment significantly attenuated the intensity of CC3 in NeuN^+^ neurons in the cerebral cortex of tMCAO mice compared with vehicle cohorts (**Fig. 7G**). To evaluate axonal damage and neuronal health, we performed double IF staining using the monoclonal antibody SMI32 for labeling injured axons and MAP2a for visualizing neuronal cell bodies and dendrites (**Fig. 7H**). Our quantitative results showed that SMI32^+^ signal intensity was significantly mitigated in the cortex of tMCAO mice treated with ARN1468 versus vehicle (**Fig. 7I**) whereas MAP2a intensity was remarkably increased at day 7 post-tMCAO (**Fig. 7J**), suggesting that SERPINA3N inhibition protects neurons and axons from oxidative stress-induced damage. Collectively, our findings unravel a neurotoxic role of SERPINA3N in cerebral ischemia.

**Figure 7.**
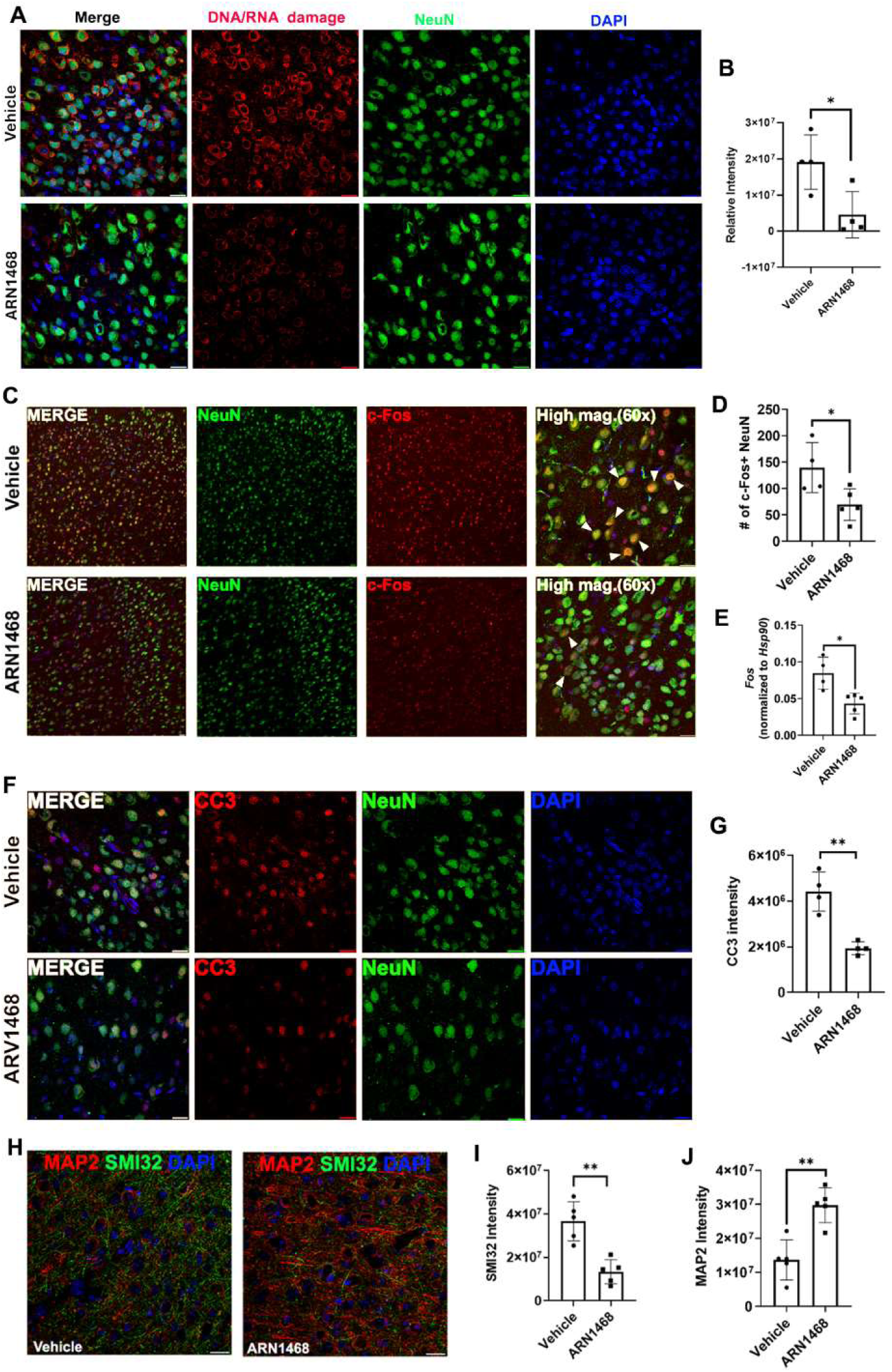
– SERPINA3N inhibition protects neurons from oxidative stress-induced damage. **(A-B)** Representative immunofluorescence images and quantification of neuronal DNA/RNA damages at Day 7 following cerebral ischemia. Sections were stained for NeuN (neuronal marker, green) and 8-OHdG, 8-OHG, and 8- oxoG (DNA/RNA damage markers, red) with DAPI (blue). (Unpaired Student’s t-test, t(df), and p value; DNA/RNA damage, t(6)=2.951, p=0.0256; n=4 per group). Scale bar, 20 µm. 60x magnification. **(C-D)** Representative immunofluorescence images and quantification of neuronal activation (c-Fos, red; NeuN, green; DAPI, blue) and **(E)** RT-qPCR analysis of Fos mRNA level at Day 7 post-cerebral ischemia. (Unpaired Student’s t- test, t(df), and p value; c-Fos, t(7)=2.715, p=0.030; Fos mRNA, t(7)=3.482, p=0.0102; n=4 Vehicle, n=5 ARN1468). Scale bar, 20 µm. 20x magnification. **(F-G)** Representative immunofluorescence images and quantification of apoptotic neurons at Day 7 post-cerebral ischemia. Sections were stained for cleaved caspase 3 (CC3, red) and NeuN (green), with DAPI (blue) counterstaining. (Unpaired Student’s t-test, t(df), and p value; CC3, t(6)=5.519, p=0.015; n=4 per group). Scale bar, 20 µm. 60x magnification. **(H-J)** Representative immunofluorescence images and quantification of neuronal degeneration. Sections were stained for MAP2 (dendritic marker, red) and SMI32 (axonal damage, green), with DAPI (blue) counterstaining. (Unpaired Student’s t-test, t(df), and p value; SMI32, t(8)=4.893, p=0.0012; MAP2, t(8)=4.591, p=0.0018; n=5 per group). Scale bar, 20 µm. 60x magnification.

## Discussion

SERPINA3N/SERPINA3 are injury-induced proteins that have been reported to be dysregulated in many neurological diseases and injuries ^4^. In acute stroke patients, higher SERPINA3 levels are significantly associated with worse disease burden and poorer neurological recovery^11,14^, highlighting the clinical importance of SERPINA3 in regulating brain damage. Yet, little is known about the expression and cellular identity of SERPINA3 in the brain in stroke patients. The tMCAO model has been used to study the expression and function of SERPINA3N in mice. In the current study, we present convincing data demonstrating that oligodendrocytes are the primary cellular origin of SERPINA3N in the brains of tMCAO mice and stroke patients. We found that antagonizing SERPINA3N function attenuates BBB disruption, reduces neuroinflammation, and mitigates ischemia-elicited neuronal stress and apoptosis in tMCAO mice. Our findings suggest that SERPINA3N is neurotoxic in cerebral ischemia. Future therapeutic designs should consider inhibiting SERPINA3N to yield beneficial effects on ischemic stroke.

The cellular origin(s) of SERPINA3N in tMCAO rodents is highly controversial. There are at least two major reasons for the controversy. First, SERPINA3N is a secreted protein. Therefore, immunoreactive localization of SERPINA3N in a cell population does not necessarily reflect the cellular origins. Second, Serpina3n mRNA in a cell population does not necessarily translate into protein. We have recently shown that Serpina3n mRNA transcripts are abundant in astrocytes during CNS demyelination but they do not bind to ribosomes and translate into SERPINA3N protein^15^. To overcome these two technical uncertainties, we employed Serpina3n- tdTom reporter mice ^15^ to define the cellular origin(s) of SERPINA3N after cerebral ischemia. Our results demonstrate that tdTom is absent from the brain subjected to sham surgery or contralateral brain hemisphere to the ischemic injury side but remarkably increased in the ipsilateral hemisphere, suggesting that SERPINA3N is an injury-induced molecule. We found that neurons did not express tdTom throughout the post-tMCAO time course we assessed. This finding is in sharp contrast to previous studies reporting that neurons are the primary cellular source of SERPINA3N^10,13^. Our conclusion is strengthened by our meta-analysis of single cell transcriptomics (**Fig. 3**) demonstrating minimal levels of Serpina3n mRNA in tMCAO brains. Interestingly, we found that damaged neurons in the acute phase of tMCAO displayed intense autofluorescence of SERPINA3N and NeuN co-localization (**Fig. S3A**) and that they were negative for tdTom (**Fig. S3B**). It is possible that SERPINA3N immunoreactive signal observed in previous studies^10,13^ may result from autofluorescence or deposition from other non-neuronal cells. Different from previous reports^13^, activated microglia/macrophages were negative for Serpina3n-tdTom (**Fig. S2**), suggesting a neglectable contribution of brain SERPINA3N by myeloid cells. We noticed that oligodendrocyte progenitor cells (OPCs) have high levels of Serpina3n mRNA transcripts assessed by single cell transcriptomics (**Fig. 3D**). However, it seems that these transcripts do not translate into SERPINA3N protein given the absence of tdTom in OPCs (**Fig. S4**). These results highlight the importance of using reporter mice to define the cellular origin of SERPINA3N particularly in the damaged/injured brain.

In this study, we identified oligodendrocytes as the major source of SERPINA3N in the cortex, white matter, and striatum of ischemic brain, particularly at late stages day 7 post-tMCAO. By day 7 post-tMCAO, greater than 80% of tdTom^+^ cells are oligodendrocytes in the cortex and corpus callosum of ischemic brain. Consistent with our tMCAO data, we found that oligodendrocytes were the major cellular source of SERPINA3 followed by astrocytes in human ischemic stroke patient tissues. Astrocytes were the second major cellular sources of SERPINA3N during the intermediate phases of tMCAO by day 3 post-tMCAO (**Fig. 2D**), particularly in the cerebral cortex where astrocytes are abundant. Within different populations of oligodendrocytes, Serpina3n is primarily enriched in the final developmental stages, i.e. myelinating OLs (cluster 6) and mature OLs (cluster 8) (**Fig. 3E-F**). Functional enrichment assay indicates that Serpina3n-expressing OLs (cluster 6 and 8) upregulate biological processes of stress responses such as cellular respiration, apoptotic signaling pathway, and inflammatory response, and downregulate those of normal oligodendroglia functions such as focal adhesion assembly, lipid synthesis, Rho signal transduction (**Fig. 3G-H**). These results suggest that OLs may not be passive responders or victims to cerebral ischemia but that they may play essential active roles in regulating ischemic brain damage through expressing and secreting SERPINA3N.

The function of SERPINA3N in ischemic brain damage remains elusive. Correlation studies of stroke patients ^11,14^ suggest that SERPINA3 may play a detrimental role in stroke brain pathophysiology. Using tMCAO rodent models, previous studies reported that neuron-derived SERPINA3N was an endogenous protective mechanism against BBB damage^13^ and that astrocyte/neuron-derived SERPINA3N attenuate brain damage by reducing neuronal apoptosis and neuroinflammation^10^. These conclusions are conceptually incompatible with the correlation studies of stroke patients^11,14^. Furthermore, recent studies demonstrated that reactive astrocyte-derived SERPINA3N induced BBB disruption in organoid cultures ^37^. More recently, we reported that oligodendrocyte-derived SERPINA3N promoted neuroinflammation and glial activation in the demyelinating and normal aging CNS^15^. From a therapeutic perspective, it is important and necessary to define the effect of pharmacologically manipulating SERINA3N on ischemic brain damage irrespective of its cellular sources. We used ARN1468 ^19^ to antagonize SERPINA3N’s function ^38^ and study the therapeutic values in intervening brain damage after tMCAO.

Different from previous studies reporting a protective role of SERPINA3N in tMCAO, we conclude that SERPINA3N induction represents a neurotoxic mechanism perpetuating brain pathophysiology after ischemic injury. Specifically, SERPINA3N antagonism significantly reduces BBB permeability and augments the expression of tight junction proteins that are required for BBB functional integrity. Accordingly, immune cell infiltration into the brain is reduced in tMCAO mice treated by ARN1468. In line with the conclusions from recent studies^15,37^, antagonizing SERPINA3N reduces pro-inflammatory response of the brain to ischemic injury after tMCAO and diminishes microglial/astroglial activation as well. Impressively, SERPINA3N inhibition is protective from ischemic injury-induced neuronal stress and apoptosis. Given these beneficial effects, ARN1468 treatment improves functional recovery of tMCAO mice.

The reasons for the different conclusions between previous studies^10,13^ and the current one remain elusive, yet several possibilities may interpret the difference. Technically, the duration of the middle cerebral artery occlusion (MCAO) is very critical in determining the rodent survival rate and brain pathological severity. We found that 1-hour MCAO used in prior studies ^10,13^ resulted in >30% mortality rate within 3 days post-tMCAO. Instead, we used 45 min occlusion time to reduce brain damage severity and increase survival rate (>90% throughout the disease time course), which mimics clinical scenarios of ischemic stroke patients. It is possible that SERPINA3N’s function may depend on brain injury severity. Second, we treated mice after tMCAO and tested the therapeutic potential of SERPINA3N in reducing ischemic brain injury whereas prior studies used AAV-mediated knockdown or overexpression 3-4 weeks prior to tMCAO and tested the prophylactic effect of SERPIA3N on brain susceptibility to ischemic brain injury. Third, neurons do not express SERPINA3N in tMCAO mice (Fig. 2, Fig. S3, Fig. 3D) and stroke patient (Fig. 4B). It is possible that the prophylactic effect, if any, of neuron-specific SERPINA3N knockdown ^13^ may be attributed to SERPINA3N from non-neuronal sources. Fourth, it is possible that SERPINA3N secreted from oligodendrocytes may be functionally different in regulating ischemic brain damage from that secreted from astrocytes or cells in the peripheral organs such as liver which highly upregulate SERPINA3N upon injury^4^. Future studies using cell-specific tools of Serpina3n knockout or overexpression are needed to answer these questions.

In summary, using reporter mice our study resolves a longstanding controversy regarding the cellular origin of SERPINA3N and concludes that oligodendrocytes but not neurons are the major source of brain SERPINA3N after ischemic brain damage. We demonstrate that SERPINA3N antagonism is a viable therapeutic option for functional recovery in ischemic stroke mice through mechanisms involving BBB function preservation, inflammation reduction, and neuronal protection.

### Limitations of the study

First, although ARN1468 has been shown to specifically inhibit SERPINA3N function^19,38^, potential off-target effects cannot be excluded. Second, pharmacological inhibition is very informative in testing the therapeutic value of SERPINA3N in ischemic stroke, the precise role of oligodendrocyte- versus astrocyte-derived SERPINA3N cannot be defined. From a basic biological perspective, it is important to tease out the cell-specific role of SERPINA3N in regulating brain pathophysiology in tMCAO mice and stroke patients. Third, the molecular mechanisms downstream of SERPINA3N in regulating stroke pathophysiological remain elusive. It is possible that serine protease-dependent and/or independent functions of SERPINA3N may play a role. Fourth, the study assessed mice up to 7 days post-tMCAO, the long-term impact of SERPINA3N inhibition on chronic tissue remodeling, remyelination, and functional recovery warrants further investigation. The genetic tools of cell-specific conditional knockout ^15^ and conditional overexpression^39^ will help overcome the limitations. Further drug screen identifying SERPINA3N specific inhibitors without any off-target will promote the translation of SERPINA3N into clinical trials.

## Materials and Methods

### Animals

C57BL/6 mice (8-12 weeks old, weight 20-25 g, Male) were used in this study. All animal experiments were conducted with University of California Davis Institutional Animal Care and Use Committee (IACUC) approval, and in accordance with the US Public Health Service’s Policy on Humane Care and Use of Laboratory Animals. All mice were housed in 12-h light/dark cycle, temperature of 20-22 °C, and humidity of 50-60 % environment, with access to food and water ad libitum.

The SERPINA3N-tdTomato reporter mice were generated using CRISPR/Cas9-based EGE technology. Heterozygote SERPINA3N-tdTomato^+/-^ mice were used by crossing C57BL/6J background wild-type mice purchased from the Jackson Laboratory.

### Transient middle cerebral artery occlusion (tMCAO)

To establish large vessel ischemia, tMCAO was performed as previously described^40^. Briefly, mice were deeply anesthetized with a mixture of ketamine (100 mg/kg) and xylazine (10 mg/kg). In brief, the neck skin was incised along the middle line, and right common carotid artery (rCCA) was exposed. The lower portion of the rCCA and the external carotid artery were permanently ligated with 6-0 silk sutures. The upper portion of the rCCA was temporarily ligated for making a small hole in the middle of the rCCA. A silicon-coated MCAO filament (Cat#6021910, Doccol Inc.) was then introduced into the MCA via circle of Willis. After 45 mins of occlusion, the filament was withdrawn to restore blood flow. For sham surgery, all procedures were identical to those in the tMCAO group except for the occlusion step.

### Treatment

As SERPINA3N inhibitor, ARN1468^19^ (MCE, 5 mg/kg) was injected twice daily for 7 days following tMCAO. As a control, vehicle (DMSO, Sigma-Aldrich, D2438) was injected intraperitoneally at the same time points.

### Immunohistochemistry (IHC)

For histological studies, mice were deeply anesthetized with a mixture of ketamine/xylazine and transcardially perfused with cold phosphate-buffered saline (PBS). After brain harvest, cerebra dissected coronally, and samples from each mouse were immediately placed on dry ice for subsequent protein or RNA extraction or fixed in 4% paraformaldehyde (PFA, Electron Microscopy Science), followed by cryoprotected in 30% sucrose, and embedded in optimal cutting temperature compound prior to cryostat sectioning. Cerebral cryostat sections were cut into 14 µm-thick for immunofluorescence. The sections were incubated with rabbit anti-RFP preabsorbed (RRID: AB_2209751, 1:200, Rockland, 600-401-379) to amplify endogenous tdTomato signal with each cell type-specific antibodies; rabbit anti-SERPINA3 (RRID: AB_1079361, 1:100, Atlas Antibodies, HPA002560), goat anti-Serpina3n (RRID: AB_2270116, 1:200, R&D system, AF4709), mouse anti-NeuN (RRID: AB_2298772, 1:200, Millipore, MAB377), mouse anti-GFAP (RRID: AB_11212597, 1:200, Millipore, MAB360), goat anti-GST-π (RRID: AB_2728849, 1:200, Abcam, ab53943), rat anti-CD68 (RRID: AB_322219, 1:200, Bio-Rad, MCA1957), rabbit anti-CD31 (RRID: AB_2757159, 1:100, Abclonal, A0378), chicken anti- MAP2 (RRID: AB_2138153, 1:200, Abcam, Ab5392), mouse anti-SMI32 (RRID: AB_2564642, 1:200, BioLegend), 801701), mouse anti-claudin 5 (RRID: AB_2533200, 1:50, Thermo Fisher Scientific, 35-2500), mouse anti-DNA/RNA damage (RRID: AB_940049, 1:100, Abcam, Ab62623), rabbit anti-CC3 (RRID: AB_2341188, 1:200, Cell Signaling, 9661), rat anti-Ly6C/G (RRID: AB_467730, 1:200, Invitrogen, 14-5931-82), hamster anti-CD3e (RRID: AB_394727, 1:200, BD, 553238), rabbit anti-c-Fos (RRID: AB_2629503, 1:100, Santa Cruz, Sc-52) then incubated with fluorescently tagged secondary antibodies, counterstained with 4’,6’-diamidino-2- phenylindole (DAPI, 1:1000, Thermo Fisher Scientific, PI62247). All confocal images were obtained from a Nikon A1 laser scanning confocal microscope (Nikon, Tokyo, Japan). Images across treatment groups were analyzed with the same intensity thresholds.

### Protein extraction and Western blot analysis

Brain tissues were homogenized using RIPA buffer (Thermo Fisher Scientific, 37585) and the total amount of protein was quantified by BCA assay kit (Thermo Fisher Scientific, 23227). The proteins were separated using SDS-PAGE gel electrophoresis and transferred to PVDF membranes. Membranes then were blocked with the blocking buffer (Bio-Rad, 12010020) for 10 mins at room temperature, followed by primary antibodies; goat-anti SERPINA3N (1:1000, RRID: AB_355395, R&D System, AF4709) and Rabbit-anti beta I tubulin(1:2000, RRID: AB_880625, Abcam, ab40742), and then with HRP-conjugated secondary antibodies. The membranes were developed to detect band intensities using a gel imaging system (UVP GelStudio Plus, Analytik Jena).

### Assessment of blood-brain barrier leakage by Evans Blue (EB)

To assess the BBB integrity, 200 µL of 0.5% EB solution (M4159, Sigma-Aldrich) was injected via retro-orbital space on day 1 and 7 following tMCAO, respectively. The procedures were followed by previously described^41^. After allowing EB dye to circulate vessels for 30 mins, mice were subjected to transcardial perfusion with ice-cold PBS to completely drain the circulating EB. Quantitative analyses were conducted as previously described^42^. The final EB absorbance was measured at 610 nM using a microplate reader (Accuris MR9600).

### RNA extraction and real-time quantitative PCR (RT-qPCR)

RNA was isolated from the brain tissues using AFTSpin Tissue Fast RNA extraction kit (ABclonal, RK30120). The final concentration of RNA was determined using a spectrophotometer (Thermo Fisher Scientific, Nanodrop 2000). cDNA was synthesized using the ABScript II cDNA First-Strand Synthesis Kit (ABclonal, RK20400). The The RT-qPCR analyses were conducted with the 2x Universal SYBR Green Fast qPCR Mix (ABclonal, RM21203) on a thermocycler system (qTOWERiris, Analytik Jena, Germany). mRNA expression levels were normalized to the housekeeping gene Hsp90. Relative expression was calculated using the 2^(-ΔCt) method, method -ΔCt was defined as the difference between the Ct values of the target gene and Hsp90. Expression in control samples was set to 1 for comparison. The primer sequences used in this study are provided in **supplementary table 1**.

### Behavioral tests

To evaluate the neurological deficit, modified neurological severity score (mNSS) testing was performed by blind observers for 7 days as previously described^43^. The mNSS evaluates neurological function across motor, sensory, balance, and reflex domains, and also accounts for abnormal behaviors such as seizures. Scoring ranges from 0 to 18, where 0 indicates normal function and 18 reflects the most severe neurological impairments.

### Human samples

For the human study, anonymized formalin-fixed paraffin-embedded brain samples including the core and peri-infarct region from autopsies of patients with ischemic stroke (n=4) were obtained from the archives of the Palo Alto Veterans Affairs Health Care System, Palo Alto, California. For each case, 8 µm-thick sections were stained with hematoxylin/eosin and labeled for each cell type-specific marker and anti-SERPINA3 by immunofluorescence using antibodies to; neurons (NeuN, RRID: AB_2298772, Cat#MAB377), astrocytes (GFAP, RRID: AB_1212597, Cat#MAB360, Millipore), oligodendrocytes (GST-pi, RRID: AB_2279558, Cell Signaling, Cat#3369), microglia/macrophages (CD68, RRID: AB_3100585, BioRad, Cat#MCA1957), and SERPINA3 (RRID: AB_10344682, Proteintech, Cat#12192-1-AP).

### Statistics

Statistical analyses were performed using GraphPad Prism (version 8, GraphPad Software Inc., USA). Normality of data distribution was assessed using the Shapiro-Wilk test. Data are presented as mean ± SD unless otherwise specified. For comparisons between two groups (Vehicle vs ARN1468), a two-tailed-tailed Student’s *t-*test was used. For multiple comparisons, among three or more groups, one-way ANOVA followed by Tukey’s multiple comparisons test was applied. Statistical significance was set as *p* < 0.05.

## Acknowledgement

We thank the funding agencies of NIH (R01NS123080, R01NS123165, R01NS134887 to FG) and Shriners Hospitals for Children (85101-NCA-22, 85113-NCA-23 to FG, 84331-NCA-24 to JP) for supporting the work. We sincerely thank the patients and their families for their invaluable tissue donations, which made this study possible.

## Data availability

Data will be made available on request. Single cell transcriptomics datasets were downloaded from the Gene Expression Omnibus (GEO) database. GSE identifications were assigned as follows: GSE197731, GSE227651, GSE245386

## Author contributions

Design and Conceptualization: JP and FG; Funding Acquisition and Supervision: JP and FG; Data generation: JP, BB, and HK; Writing-original draft: JP and FG; Writing-review and editing: JP, BB, RS, MY, and FG; Figures and Tables: JP and BB; Manuscript finalization: FG. All authors read and approved of the final manuscript.

## References

1. Xu, A., Zhang, H., Zhang, Y., Wu, J., and Huang, Z. (2025). Ischemic stroke and intervention strategies based on the timeline of stroke progression: Review and prospects. Acta Pharm Sin B 15, 4543–4581. 10.1016/j.apsb.2025.07.026.

2. Sekerdag, E., Solaroglu, I., and Gursoy-Ozdemir, Y. (2018). Cell Death Mechanisms in Stroke and Novel Molecular and Cellular Treatment Options. Curr Neuropharmacol 16, 1396–1415. 10.2174/1570159X16666180302115544.

3. Horvath, A.J., Irving, J.A., Rossjohn, J., Law, R.H., Bottomley, S.P., Quinsey, N.S., Pike, R.N., Coughlin, P.B., and Whisstock, J.C. (2005). The murine orthologue of human antichymotrypsin: a structural paradigm for clade A3 serpins. J Biol Chem 280, 43168–43178. 10.1074/jbc.M505598200.

4. Zhu, M., Lan, Z., Park, J., Gong, S., Wang, Y., and Guo, F. (2024). Regulation of CNS pathology by Serpina3n/SERPINA3: The knowns and the puzzles. Neuropathol Appl Neurobiol 50, e12980. 10.1111/nan.12980.

5. Fissolo, N., Matute-Blanch, C., Osman, M., Costa, C., Pinteac, R., Miro, B., Sanchez, A., Brito, V., Dujmovic, I., Voortman, M., et al. (2021). CSF SERPINA3 Levels Are Elevated in Patients With Progressive MS. Neurol Neuroimmunol Neuroinflamm 8. 10.1212/NXI.0000000000000941.

6. DeKosky, S.T., Ikonomovic, M.D., Wang, X., Farlow, M., Wisniewski, S., Lopez, O.L., Becker, J.T., Saxton, J., Klunk, W.E., Sweet, R., et al. (2003). Plasma and cerebrospinal fluid alpha1-antichymotrypsin levels in Alzheimer’s disease: correlation with cognitive impairment. Ann Neurol 53, 81–90. 10.1002/ana.10414.

7. Abraham, C.R., Selkoe, D.J., and Potter, H. (1988). Immunochemical identification of the serine protease inhibitor alpha 1-antichymotrypsin in the brain amyloid deposits of Alzheimer’s disease. Cell 52, 487–501. 10.1016/0092-8674(88)90462-x.

8. Han, X., Lei, Q., Liu, H., Zhang, T., and Gou, X. (2024). SerpinA3N Regulates the Secretory Phenotype of Mouse Senescent Astrocytes Contributing to Neurodegeneration. J Gerontol A Biol Sci Med Sci 79. 10.1093/gerona/glad278.

9. Wang, Z.M., Liu, C., Wang, Y.Y., Deng, Y.S., He, X.C., Du, H.Z., Liu, C.M., and Teng, Z.Q. (2020). SerpinA3N deficiency deteriorates impairments of learning and memory in mice following hippocampal stab injury. Cell Death Discov 6, 88. 10.1038/s41420-020-00325-8.

10. Zhang, Y., Chen, Q., Chen, D., Zhao, W., Wang, H., Yang, M., Xiang, Z., and Yuan, H. (2022). SerpinA3N attenuates ischemic stroke injury by reducing apoptosis and neuroinflammation. CNS Neurosci Ther 28, 566–579. 10.1111/cns.13776.

11. Zheng, P., Qi, Z., Gao, B., Yao, Y., Chen, J., Cong, H., Huang, Y., and Shi, F.D. (2025). SERPINA3 predicts long-term neurological outcomes and mortality in patients with intracerebral hemorrhage. Cell Death Dis 16, 218. 10.1038/s41419-025-07551-x.

12. Zamanian, J.L., Xu, L., Foo, L.C., Nouri, N., Zhou, L., Giffard, R.G., and Barres, B.A. (2012). Genomic analysis of reactive astrogliosis. J Neurosci 32, 6391–6410. 10.1523/JNEUROSCI.6221-11.2012.

13. Li, F., Zhang, Y., Li, R., Li, Y., Ding, S., Zhou, J., Huang, T., Chen, C., Lu, B., Yu, W., et al. (2023). Neuronal Serpina3n is an endogenous protector against blood brain barrier damage following cerebral ischemic stroke. J Cereb Blood Flow Metab 43, 241–257. 10.1177/0271678X221113897.

14. Hu, X., Xiao, Z.S., Shen, Y.Q., Yang, W.S., Wang, P., Li, P.Z., Wang, Z.J., Pu, M.J., Zhao, L.B., Xie, P., and Li, Q. (2023). SERPINA3: A novel inflammatory biomarker associated with cerebral small vessel disease burden in ischemic stroke. CNS Neurosci Ther. 10.1111/cns.14472.

15. Wang, Y., Zhu, M., Park, J., Kim, H., Ge, X., Chen, X., Chen, B., Ding, S., Lin, W., and Guo, F. (2026). Injury-transduced oligodendrocytes modulate neuroinflammation and glial activation in the diseased and non-diseased CNS. Nat Commun (In Press).

16. Ruan, Z., Cao, G., Qian, Y., Fu, L., Hu, J., Xu, T., Wu, Y., and Lv, Y. (2023). Single-cell RNA sequencing unveils Lrg1’s role in cerebral ischemia‒reperfusion injury by modulating various cells. J Neuroinflammation 20, 285. 10.1186/s12974-023-02941-4.

17. Zeng, F., Cao, J., Hong, Z., Liu, Y., Hao, J., Qin, Z., Zou, X., and Tao, T. (2023). Single-cell analyses reveal the dynamic functions of Itgb2(+) microglia subclusters at different stages of cerebral ischemia-reperfusion injury in transient middle cerebral occlusion mice model. Front Immunol 14, 1114663. 10.3389/fimmu.2023.1114663.

18. Kim, S., Lee, W., Jo, H., Sonn, S.K., Jeong, S.J., Seo, S., Suh, J., Jin, J., Kweon, H.Y., Kim, T.K., et al. (2022). The antioxidant enzyme Peroxiredoxin-1 controls stroke-associated microglia against acute ischemic stroke. Redox Biol 54, 102347. 10.1016/j.redox.2022.102347.

19. Colini Baldeschi, A., Zattoni, M., Vanni, S., Nikolic, L., Ferracin, C., La Sala, G., Summa, M., Bertorelli, R., Bertozzi, S.M., Giachin, G., et al. (2022). Innovative Non-PrP-Targeted Drug Strategy Designed to Enhance Prion Clearance. J Med Chem 65, 8998–9010. 10.1021/acs.jmedchem.2c00205.

20. Yu, F., Kamada, H., Niizuma, K., Endo, H., and Chan, P.H. (2008). Induction of mmp-9 expression and endothelial injury by oxidative stress after spinal cord injury. J Neurotrauma 25, 184–195. 10.1089/neu.2007.0438.

21. Yang, C., Hawkins, K.E., Dore, S., and Candelario-Jalil, E. (2019). Neuroinflammatory mechanisms of blood-brain barrier damage in ischemic stroke. Am J Physiol Cell Physiol 316, C135–C153. 10.1152/ajpcell.00136.2018.

22. Dong, X., Song, Y.N., Liu, W.G., and Guo, X.L. (2009). Mmp-9, a potential target for cerebral ischemic treatment. Curr Neuropharmacol 7, 269–275. 10.2174/157015909790031157.

23. Schadlich, I.S., Vienhues, J.H., Jander, A., Piepke, M., Magnus, T., Lambertsen, K.L., Clausen, B.H., and Gelderblom, M. (2022). Interleukin-1 Mediates Ischemic Brain Injury via Induction of IL-17A in gammadelta T Cells and CXCL1 in Astrocytes. Neuromolecular Med 24, 437–451. 10.1007/s12017-022-08709-y.

24. Jin, R., Liu, L., Zhang, S., Nanda, A., and Li, G. (2013). Role of inflammation and its mediators in acute ischemic stroke. J Cardiovasc Transl Res 6, 834–851. 10.1007/s12265-013-9508-6.

25. Fanzhou, R., Yuding, L., Pingchuan, L., Hai, X., Xu, L., and Jian, W. (2025). The role of TNF signaling pathway in post-stroke cognitive impairment: a systematic review. Ann Med 57, 2543519. 10.1080/07853890.2025.2543519.

26. Takada, Y.K., Fujita, M., and Takada, Y. (2022). Pro-Inflammatory Chemokines CCL5, CXCL12, and CX3CL1 Bind to and Activate Platelet Integrin alphaIIbbeta3 in an Allosteric Manner. Cells 11. 10.3390/cells11193059.

27. van Wageningen, T.A., Vlaar, E., Kooij, G., Jongenelen, C.A.M., Geurts, J.J.G., and van Dam, A.M. (2019). Regulation of microglial TMEM119 and P2RY12 immunoreactivity in multiple sclerosis white and grey matter lesions is dependent on their inflammatory environment. Acta Neuropathol Commun 7, 206. 10.1186/s40478-019-0850-z.

28. Clarke, L.E., Liddelow, S.A., Chakraborty, C., Munch, A.E., Heiman, M., and Barres, B.A. (2018). Normal aging induces A1-like astrocyte reactivity. Proc Natl Acad Sci U S A 115, E1896–E1905. 10.1073/pnas.1800165115.

29. Liddelow, S.A., Guttenplan, K.A., Clarke, L.E., Bennett, F.C., Bohlen, C.J., Schirmer, L., Bennett, M.L., Munch, A.E., Chung, W.S., Peterson, T.C., et al. (2017). Neurotoxic reactive astrocytes are induced by activated microglia. Nature 541, 481–487. 10.1038/nature21029.

30. Chen, H., Yoshioka, H., Kim, G.S., Jung, J.E., Okami, N., Sakata, H., Maier, C.M., Narasimhan, P., Goeders, C.E., and Chan, P.H. (2011). Oxidative stress in ischemic brain damage: mechanisms of cell death and potential molecular targets for neuroprotection. Antioxid Redox Signal 14, 1505–1517. 10.1089/ars.2010.3576.

31. Wu, L., Xiong, X., Wu, X., Ye, Y., Jian, Z., Zhi, Z., and Gu, L. (2020). Targeting Oxidative Stress and Inflammation to Prevent Ischemia-Reperfusion Injury. Front Mol Neurosci 13, 28. 10.3389/fnmol.2020.00028.

32. Manzanero, S., Santro, T., and Arumugam, T.V. (2013). Neuronal oxidative stress in acute ischemic stroke: sources and contribution to cell injury. Neurochem Int 62, 712–718. 10.1016/j.neuint.2012.11.009.

33. Jurcau, A., and Ardelean, A.I. (2022). Oxidative Stress in Ischemia/Reperfusion Injuries following Acute Ischemic Stroke. Biomedicines 10. 10.3390/biomedicines10030574.

34. Molina, P., Belda, X., Beriain, S., Serrano, S., Compte, G., Andero, R., and Armario, A. (2025). Dynamics of stress-induced c-fos expression in the rat prelimbic cortex: lessons from intronic and mature RNA and protein analyses. Neurobiol Stress 36, 100729. 10.1016/j.ynstr.2025.100729.

35. Miles, A.N., and Knuckey, N.W. (1998). Apoptotic neuronal death following cerebral ischaemia. J Clin Neurosci 5, 125–145. 10.1016/s0967-5868(98)90027-3.

36. Mao, R., Zong, N., Hu, Y., Chen, Y., and Xu, Y. (2022). Neuronal Death Mechanisms and Therapeutic Strategy in Ischemic Stroke. Neurosci Bull 38, 1229–1247. 10.1007/s12264-022-00859-0.

37. Kim, H., Leng, K., Park, J., Sorets, A.G., Kim, S., Shostak, A., Embalabala, R.J., Mlouk, K., Katdare, K.A., Rose, I.V.L., et al. (2022). Reactive astrocytes transduce inflammation in a blood-brain barrier model through a TNF-STAT3 signaling axis and secretion of alpha 1-antichymotrypsin. Nat Commun 13, 6581. 10.1038/s41467-022-34412-4.

38. Ma, K., Zhang, C., Zhang, H., An, C., Li, G., Cheng, L., Li, M., Ren, M., Bai, Y., Liu, Z., et al. (2024). High-Salt Diet Accelerates Neuron Loss and Anxiety in APP/PS1 Mice Through Serpina3n. Int J Mol Sci 25. 10.3390/ijms252111731.

39. Zhu, M., Barman, B., Guo, C., Murali, R., Joseph, J., and Guo, F. (2025). A paradigm shift of SERPINA3N in neurobehavioral development and brain injury. BioRxiv https://www.biorxiv.org/content/10.1101/2025.09.09.675167v1.

40. Park, J., Kim, J.Y., Kim, Y.R., Huang, M., Chang, J.Y., Sim, A.Y., Jung, H., Lee, W.T., Hyun, Y.M., and Lee, J.E. (2021). Reparative System Arising from CCR2(+) Monocyte Conversion Attenuates Neuroinflammation Following Ischemic Stroke. Transl Stroke Res 12, 879–893. 10.1007/s12975-020-00878-x.

41. Yardeni, T., Eckhaus, M., Morris, H.D., Huizing, M., and Hoogstraten-Miller, S. (2011). Retro-orbital injections in mice. Lab Anim (NY) 40, 155–160. 10.1038/laban0511-155.

42. Radu, M., and Chernoff, J. (2013). An in vivo assay to test blood vessel permeability. J Vis Exp, e50062. 10.3791/50062.

43. Chen, J., Li, Y., Wang, L., Zhang, Z., Lu, D., Lu, M., and Chopp, M. (2001). Therapeutic benefit of intravenous administration of bone marrow stromal cells after cerebral ischemia in rats. Stroke 32, 1005–1011. 10.1161/01.str.32.4.1005.

