## Supplemental Figures and Table 1 for "Cellular origin and therapeutic potential of SERPINA3N in ischemic stroke"

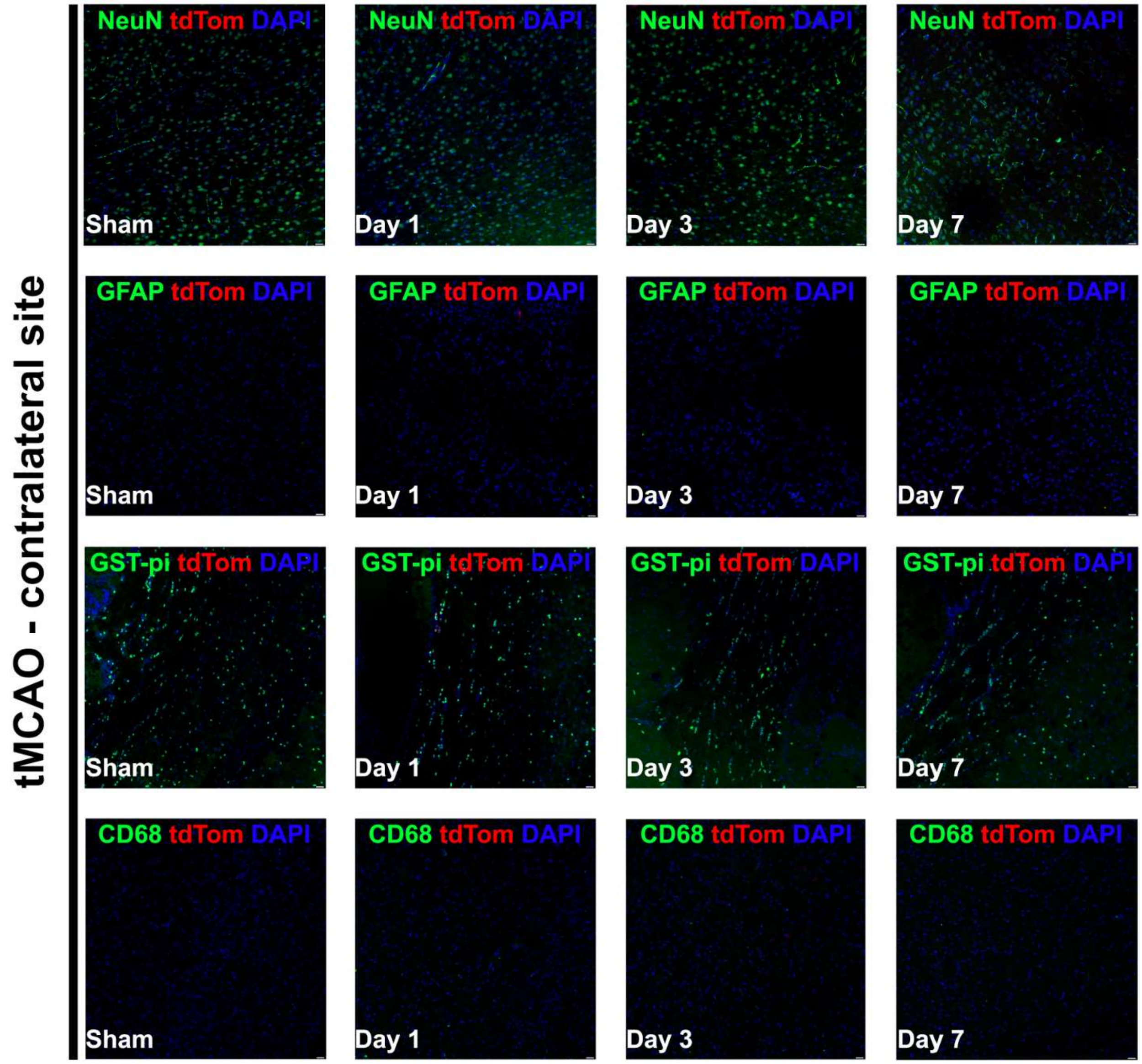


**Supplemental Figure 1 – Absence of tdTom expression in the contralateral hemisphere during cerebral ischemia.** Representative immunofluorescence images of the contralateral (non-ischemic) hemisphere at each post-tMCAO timepoint show no tdTom signal, and consequently no colocalization with cell type-specific markers, confirming that Serpina3n induction does not occur in the absence of ischemic injury. Scale bar, 20 μm; 20x magnification.

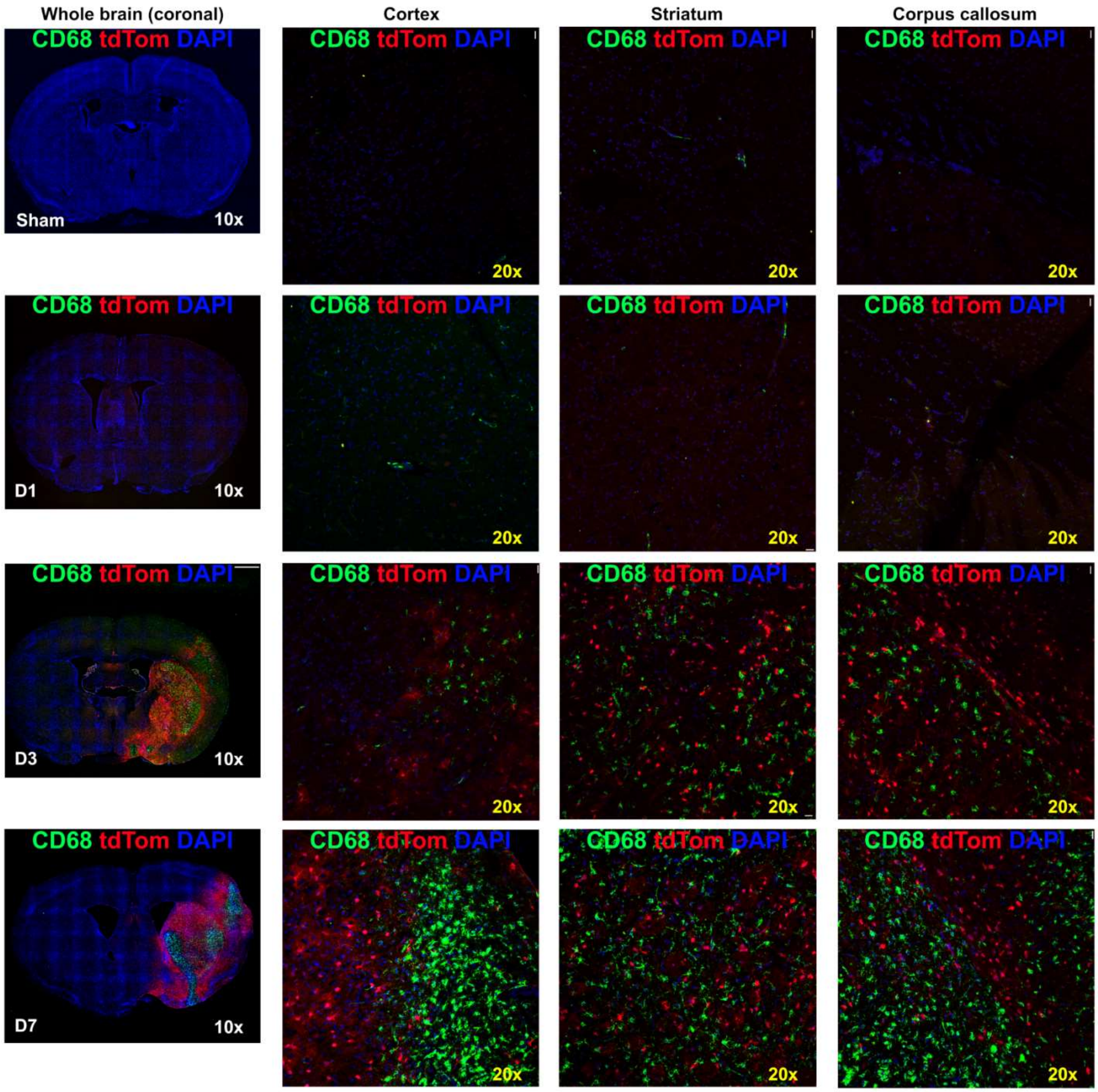


**Supplemental Figure 2 – Absence of tdTom in CD68-positive myeloid cells.** Representative whole-brain (10x) and regional (20x; cortex, striatum, corpus callosum) images in Sham and at D1, D3, and D7 post-tMCAO. CD68+ myeloid cells progressively accumulate in the ipsilateral hemisphere by D3-D7, but these cells remain tdTom-negative at all timepoints, indicating that Serpina3n inductions in not attributable to activated microglia/macrophages. Scale bar, 2mm (10x) or 20 μm (20x)

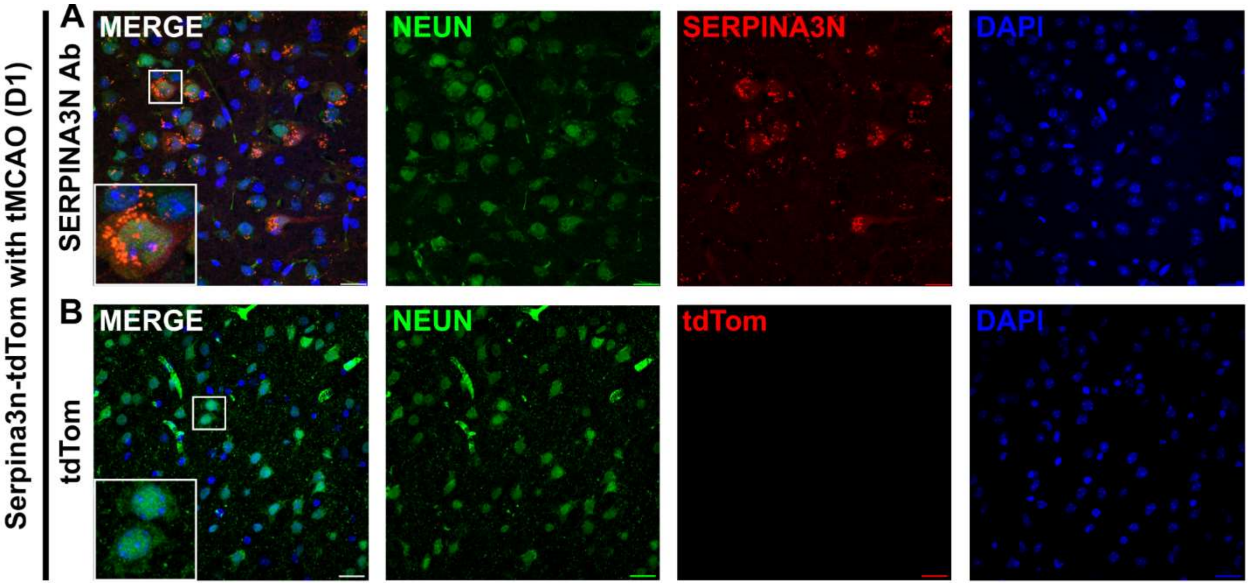


**Supplemental Figure 3 – Absence of tdTom in NeuN-positive neurons. (A)** Immunostaining with an anti-SERPINA3N antibody shows apparent signal in NeuN+ cells (inset, boxed region). **(B)** In the same Serpina3n-tdTomato reporter tissue, which directly labels cells actively transcribing Serpina3n, no tdTomato signal is detected in NeuN+ neurons (inset, boxed region). This discrepancy indicates that antibody-based immunoreactivity for SERPINA3N in neurons does not reflect genuine Seprina3n transcription, consistent with oligodendrocytes — rather than neurons — being the primary cellular source of Serpina3n after ischemic stroke. Scale bar, 20 μm. 60x magnification.

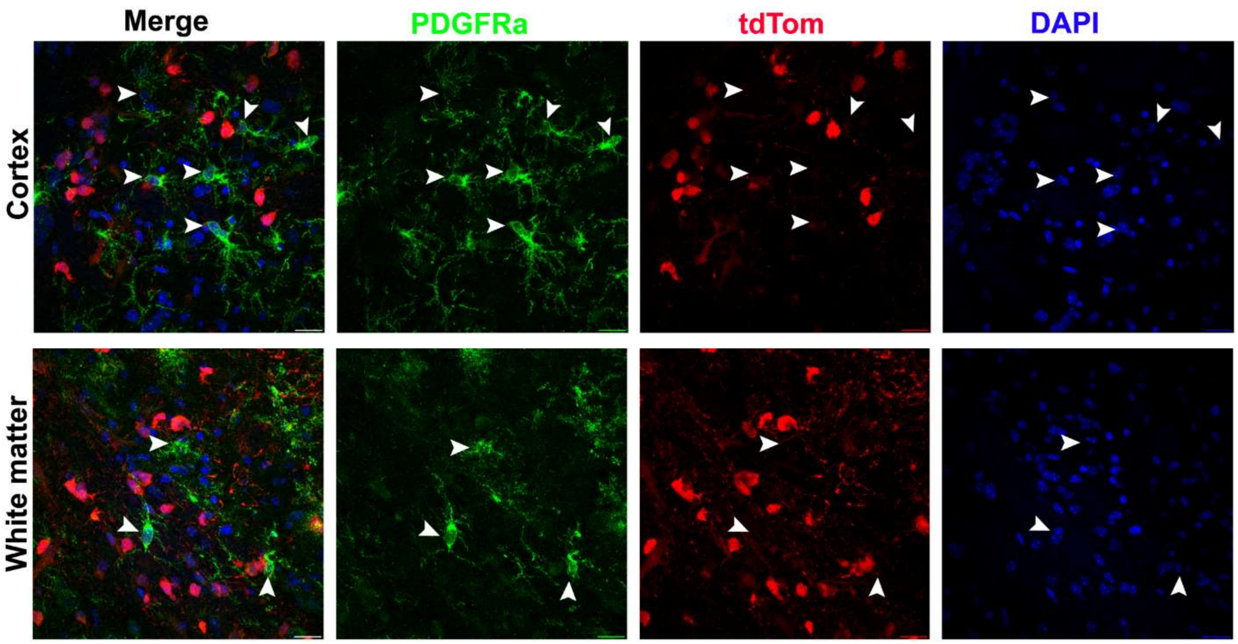


**Supplemental Figure 4 – Absence of tdTom in OPCs.** Representative immunofluorescence images of PDGFRα (green, OPC marker), tdTom (red), and DAPI (blue) in cortex and white matter. Arrowheads indicate PDGFRα+ OPCs that lack tdTom signal, demonstrating that Serpina3n is not expressed by OPCs in either region. Scale bar, 20 μm. 60x magnification.

**Supplemental Table1. RT-qPCR primer sequences**

| Gene | Forward (5′→ 3’) | Reverse (5′→ 3’) |
|---|---|---|
| ***Il1β*** | TGCCACCTTTTGACAGTGATG | TGATGTGCTGCTGCGAGATT |
| ***Tnfα*** | ACTGAACTTCGGGGTGATCG | TGATCTGAGTGTGAGGGTCTGG |
| ***Ldha*** | *TGTGGCAGACTTGGCTGAGA* | *CTGAGGAAGACATCCTCATTGATTC* |
| ***Fos*** | ATCGGCAGAAGGG GCAAAGTAG | GCAACGCAGACTT CTCATCTTCAAG |
| ***Gfap*** | GTGTCAGAAGGCCACCTCAAG | GTGTCAGAAGGCCACCTCAAG |
| ***Lcn*** | GGATGAGATTCTTGATGACACAGC | CCAGGTTCTCTGCTACAAGTCC |
| ***Ocln*** | GGACCCTGACCACTATGAAACAGACTAC | ATAGGTGGATATTCCCTGACCCAGTC |
| ***Cldn5*** | TAACCTGAAAGGGCAGCTGGAGAAAC | AGGTCCAGGCTAAGTCCTTTGGTTCAGTAG |
| ***Tjp1*** | AGCTCATAGTTCAACACAGCCTCCAG | TTCTTCCACAGCTGAAGGACTCACAG |
| ***Tmem119*** | ACTACCCATCCTCGTTCCCTGA | TAGCAGCCAGAATGTCAGCCTG |
| ***P2y12r*** | TTTCAGATCCGCAGTAAATCCAA | GGCTCCCAGTTTAGCATCACTA |
| ***Hmgb1*** | CCAAGAAGTGCTCAGAGAGGTG | GTCCTTGAACTTCTTTTTGGTCTC |
| ***Cxcl12*** | CGCCAAGGTCGTCGCCG | TTGGCTCTGGCGATGTGGC |
| ***Cd68*** | GGGGCTCTTGGGAACTACAC | GTACCGTCACAACCTCCCTG |
| ***Vim*** | TTCTCTGGCACGTCTTGACC | CTCCTGGAGGTTCTTGGCAG |
| ***Cp*** | AGGCCCTGATGAGGAACATCT | TGCTGTGAGGAGCGACCT |
| ***Lipg1*** | TACCTACACGCTGTCCTTTGGC | GCTCGCATTTCACCATCTCTGAG |
| ***Icam1*** | AACTGTGGCACCGTGCAGTC | AGGGTGAGGTCCTTGCCTACTTG |
| ***Mmp9*** | AGGGCCCCTTTCTTATTGCC | CGAGTAACGCTCTGGGGATC |
| ***Hsp90*** | AAACAAGGAGATTTTCCTCCGC | CCGTCAGGCTCTCATATCGAAT |
